# Dendritic spines are lost in clusters in Alzheimer’s disease

**DOI:** 10.1101/2020.10.20.346718

**Authors:** Mite Mijalkov, Giovanni Volpe, Isabel Fernaud-Espinosa, Javier DeFelipe, Joana B. Pereira, Paula Merino-Serrais

**Affiliations:** Department of Neurobiology, Care Sciences and Society, Karolinska Institutet, Stockholm, Sweden; Department of Physics, Goteborg University, Goteborg, Sweden; Laboratorio Cajal de Circuitos Corticales (CTB), Universidad Politécnica de Madrid, Madrid, Spain; Departamento de Neurobiología Funcional y de Sistemas, Instituto Cajal, CSIC, Madrid, Spain; Centro de Investigación Biomédica en Red sobre Enfermedades Neurodegenerativas (CIBERNED), ISCIII, Madrid, Spain; Memory Research Unit, Department of Clinical Sciences, Malmö, Lund University, Lund, Sweden

## Abstract

Alzheimer’s disease (AD) is a progressive neurodegenerative disorder characterized by a deterioration of neuronal connectivity. The pathological accumulation of tau in neurons is one of the hallmarks of AD and has been connected to the loss of dendritic spines of pyramidal cells, which are the major targets of cortical excitatory synapses and key elements in memory storage. However, the detailed mechanisms underlying the loss of dendritic spines in individuals with AD are still unclear. Here, we used graph-theory approaches to compare the distribution of dendritic spines from neurons with and without tau pathology of AD individuals. We found that the presence of tau pathology determines the loss of dendritic spines in clusters, ruling out alternative models where spine loss occurs at random locations. Since memory storage has been associated with synaptic clusters, the present results provide a new insight into the mechanisms by which tau drives synaptic damage in AD, paving the way to memory deficits through alterations of spine organization.

## INTRODUCTION

Alzheimer’s disease (AD) is the most common neurodegenerative disorder; it is characterized by a progressive loss of memory, followed by decline in other cognitive functions and ultimately dementia [1]. These clinical symptoms are thought to be caused by the underlying pathological processes associated with AD, in particular, the aggregation of tau into neurofibrillary tangles and the loss of crucial structures for synaptic communication such as dendritic spines (for simplicity, spines) [2]. However, the mechanisms that link tau pathology to spine loss in AD individuals are currently unclear. Clarifying their relation is nevertheless important to improve our understanding of the pathophysiology of AD and to develop future treatment strategies that target more specifically these mechanisms. Amongst the wide range of neural cells forming the human brain, pyramidal neurons are particularly important because they are the most abundant neurons and connect different areas of the neocortex, the hippocampus and the amygdala [3]. Previous studies have shown changes in the spine density of pyramidal neurons, affecting the number of excitatory inputs that these neurons receive and, consequently, the underlying cognitive processes they support, including memory and learning [3–6]. Moreover, there is evidence that the spines of pyramidal neurons are not uniformly distributed along their dendrites and can undergo specific plastic changes in their size and spatial location [7]. In particular, the activity of one spine can modulate the plasticity of neighboring spines through the mutual sharing of plasticity-related proteins or through the activation of synchronized synaptic inputs [8]. These changes occur across different time scales and typically result in the spatial organization of spines into groups or clusters. This phenomenon is known as the “clustered plasticity hypothesis” [9–12] and suggests that spines do not act as single functional units but are part of a complex network that organizes spines in groups to optimize the connectivity patterns between dendrites and surrounding axons. To this date, no studies have assessed how spine organization is affected by tau pathology in human individuals. This is probably due to the technical difficulties associated with studying the microanatomy of human pyramidal cells. In particular, brain tissue from deceased individuals need to be obtained with very short postmortem delays (less than 5 hours) and the intracellular injections need to be performed within a time window of 24–48 h after fixation [13]. As a further difficulty, not all dendrites are viable for further spine analysis because dendrites from neurons in advanced stages of tau pathology show virtually no spines [13, 14] and those dendrites in contact with amyloid-beta plaques display severe morphological alterations [15–17].

In this study, we overcame these issues by intracellularly injecting neurons within a short time window post mortem and by excluding those in advanced tau stages or in contact with plaques. This permits us to identify 6 neurons with 11 dendrites and 4204 spines for further analysis obtained from the hippocampal subfield *Cornu Ammonis* 1 (CA1) of an AD individual. These neurons were subsequently divided into those with intermediate levels of phospho-tau aggregates in the soma and proximal processes (SomaTau+) and those without phospho-tau aggregates in the soma nor proximal processes (SomaTau-). Moreover, we have previously described that some CA1 Lucifer yellow-injected neurons that showed phospho-tau in the soma and proximal processes (SomaTau+ neurons) also display phospho-tau in the distal portion of the dendrites in short segments of approximately 10-20 μm. In these distal dendritic regions, the dendritic spine density was lower [13]. The possible presence of phospho-tau in the distal segments of the dendrites included in the analysis has been analyzed (Supplementary Table 1 and Supplementary Table 3). In addition, no phospho-tau has been found in the distal segment of the SomaTau- dendrites included in the analysis.

Building on previous evidence showing that spines are organized in groups [18], we hypothesized that the pathological changes occurring in AD do not target single spines at random locations along the dendrite, but instead damage clusters of spines that are close to each other. To test this hypothesis, we used an approach based on graph theory, where we modeled each dendrite as an ensemble of nodes connected by edges, where the nodes correspond to spines and the edges represent physical closeness between them (i.e., the inverse distance between the spines along the dendrite). We then compared SomaTau+ to SomaTau- dendrites within the same individual to identify differences related to tau pathology without confounding factors such as the high inter-individual heterogeneity regarding both clinical and pathological characteristics as well as in pyramidal cell morphology. The analysis of the possible differences of SomaTau+ and SomaTau- dendrites between the three individuals was not performed. Our findings showed that SomaTau+ dendrites featured smaller and shorter communities of spines, which were associated with a loss of spines. In addition, using a series of simulations, we observed that these changes were not due to a random loss of spines but instead could be explained by a loss of spines at clustered locations. We have also replicated these findings in two additional independent postmortem AD individuals. Altogether, these results provide an important insight into the mechanisms underlying synaptic degeneration in AD by showing that spines are lost in clusters in AD individuals.

## RESULTS

### Classification of SomaTau+ and SomaTau- dendrites

We first analyzed the 4204 reconstructed spines from hippocampal CA1 neurons (Supplementary Table S1) in one individual with AD (P13). These neurons were injected with LY fluorescent dye and immunostained with AT8 (phospho-tau_AT8_) and PHF-1 tau (phospho-tau_PHF-1_) antibodies [13, 19]; in order to detect phospho-tau aggregations indicative of tau pathology [20]. The dendrites from neurons immunostained by either phospho-tau_AT8_ or phospho-tau_PHF-1_ in the soma were classified as SomaTau+ dendrites, whereas those not immunostained by either of these antibodies were classified as SomaTau- dendrites. Figure 1 shows examples of a Tau- neuron (Figs. 1A-C) and a Tau+ neuron (Figs. 1D-F) with zoomed-in views of their dendrites and spines (Figs. 1G-J). A high-resolution segment of one dendrite is depicted (Fig. 1K) with the red points indicating the measurement of number of spines on the dendrite axis (Fig. 1L) and the coordinates that are exported to build the graph representations of the same dendrite (Fig. 1M). In total, 11 dendrites are examined in the present study, of which 5 are SomaTau- with a total of 2266 spines, and 6 are SomaTau+ with a total of 1938 spines.

**Fig 1.**
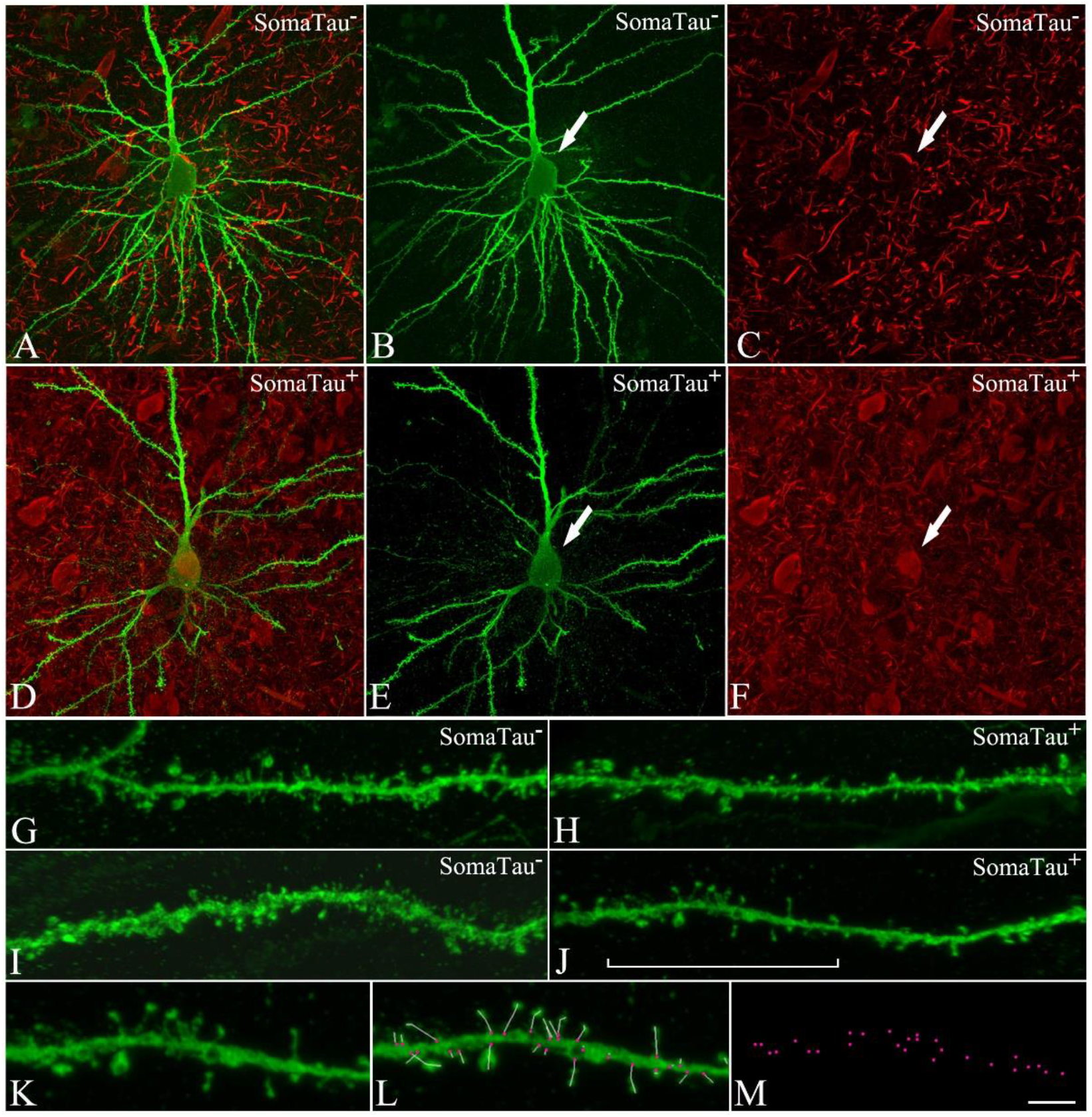
CA1 pyramidal neurons with and without tau pathology. Confocal microscopy pictures obtained after combining the channels acquired separately for Lucifer yellow (green) and phospho-tau AT8 (red), showing (**A-F**) neurons and (**G-J**) their basal dendrites, (**A-C, G-H**) with a soma free of phospho-tau AT8 (SomaTau-) and (**D-F, I-J**) with phospho-tau AT8 in an intermediate stage of neurofibrillary pathology (SomaTau+). The position of the soma is indicated with an arrow in B-C and E-F. (**K**) High resolution image of a dendritic segment indicated with a bracket in J. (**L**) The same representative image as in K showing all the spines along the dendrite marked with a white line and pink dots for their insertion points. (**M**) 3D spatial distribution of all spines insertion points. Scale bar shown in M indicates 12 μm in A-F, 5 μm in G-J, and 3 μm in K-M.

### Tau pathology is associated with smaller and shorter communities of spines

To evaluate whether the spines form closely connected communities, we modelled each dendrite as a graph, i.e., a group of nodes connected by edges. In this graph, the spines are the nodes and the edges are the connection strength between the spines. In Fig. 2, we show a tree-dimensional visual representation of the spines along a dendrite (Fig. 2A) and how we calculated the connection strength between the spines, which is proportional to their physical closeness and can be measured as the inverse distance between the spines along the dendrite (Fig. 2B). The resulting graph can be represented as an adjacency matrix (Fig. 2C), where each element in the matrix represents the connection strength between the nodes in the corresponding row and column.

**Fig 2.**
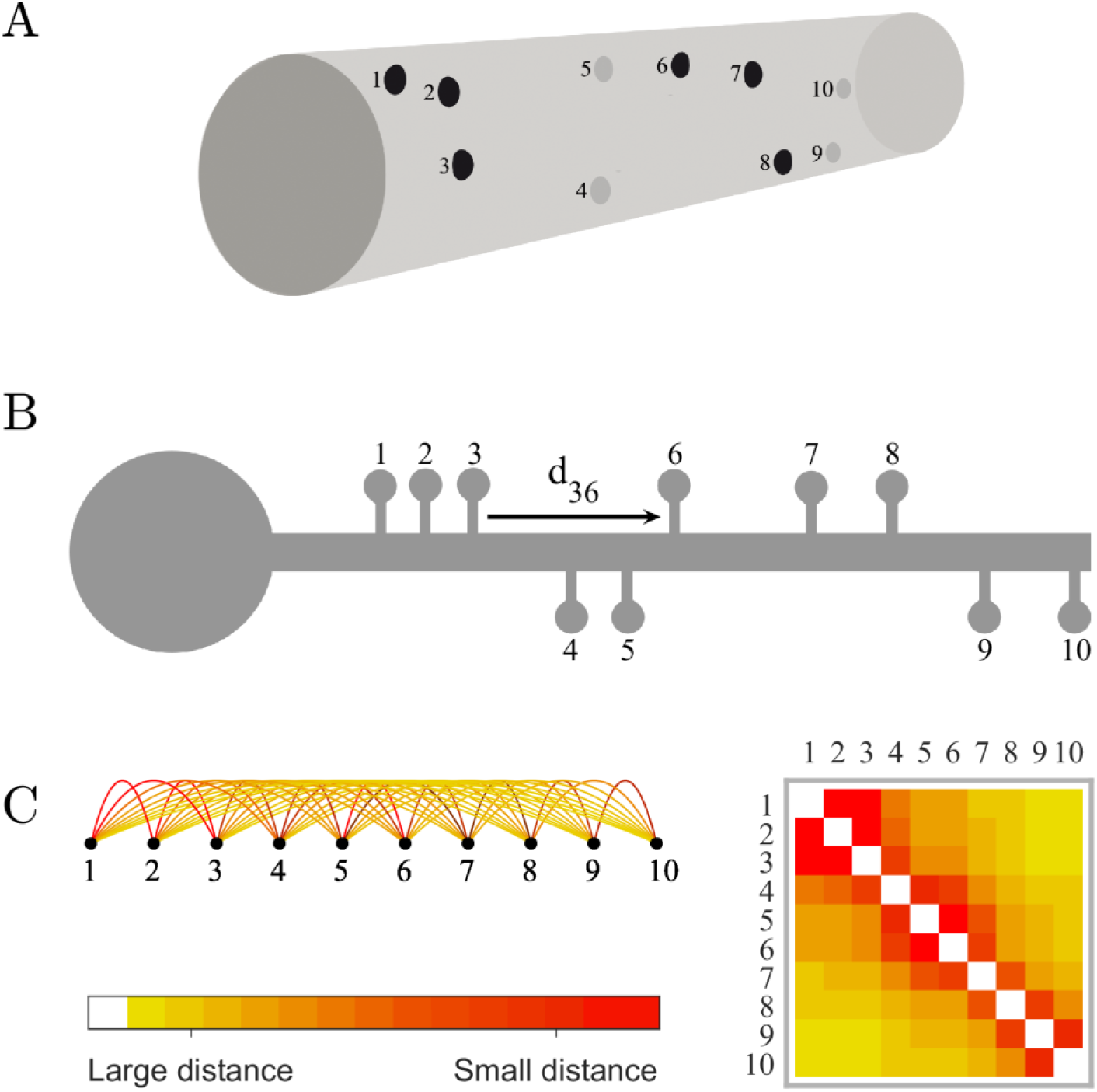
Dendrites as graphs. (**A**) 3D Visual representation of the spines (1 to 10) along a dendritic segment. (**B**) Calculation of the connection strength between pairs of spines, which is proportional to their physical closeness and can be measured as the inverse distance between the spines along the dendrite (e.g., d_36_ is the distance between spines 3 and 6, and the respective connection strength is 1/d_36_). (**C**) The resulting graph can be represented as an adjacency matrix, where each element in the matrix represents the strength of the connection between the nodes in the corresponding row and column.

We define a community as a group of spines that are tightly connected with each other and poorly connected with other groups of spines along the dendrite. Therefore, we calculate the communities by dividing the dendrite into sets of spines such that the spines belonging to each set are nearer to each other compared to other spines on the dendrite. The communities are defined using the Louvain algorithm [21]. Since this algorithm has an inherent randomness, the results are averaged over 100 community structure calculations.

In Fig. 3A we show a visual representation of an example of spine communities on a dendrite, which are visually separated into three distinct color-coded communities. We also show the corresponding adjacency matrix, where the three communities can be identified as blocks of connections. In Fig. 3B, we show the community structure for one of the SomaTau- dendrites assessed in the current study. The community structure of each dendrite can be characterized by calculating the characteristic spread between the spines within the community (characteristic community extension, CCE, shown in Fig. 3C) and the average number of spines in each community (community size, shown in Fig. 3E). This analysis shows that the characteristic extension as well as the size of the communities are significantly smaller in SomaTau+ compared to SomaTau- dendrites (*CCE*_SomaTau+_ = 17.29 ± 3.00 μm, *CCE*_SomaTau-_= 23.53 ± 3.53 μm, p < 0.001, Fig. 3D; community size: *N*_SomaTau+_ = 43.40 ± 10.72 spines, *N*_SomaTau-_= 62.40 ± 15.42 spines, p < 0.001, Fig. 3F), suggesting that the spines of SomaTau+ dendrites are organized in smaller and more tightly packed communities compared to those of the SomaTau- dendrites.

**Fig 3.**
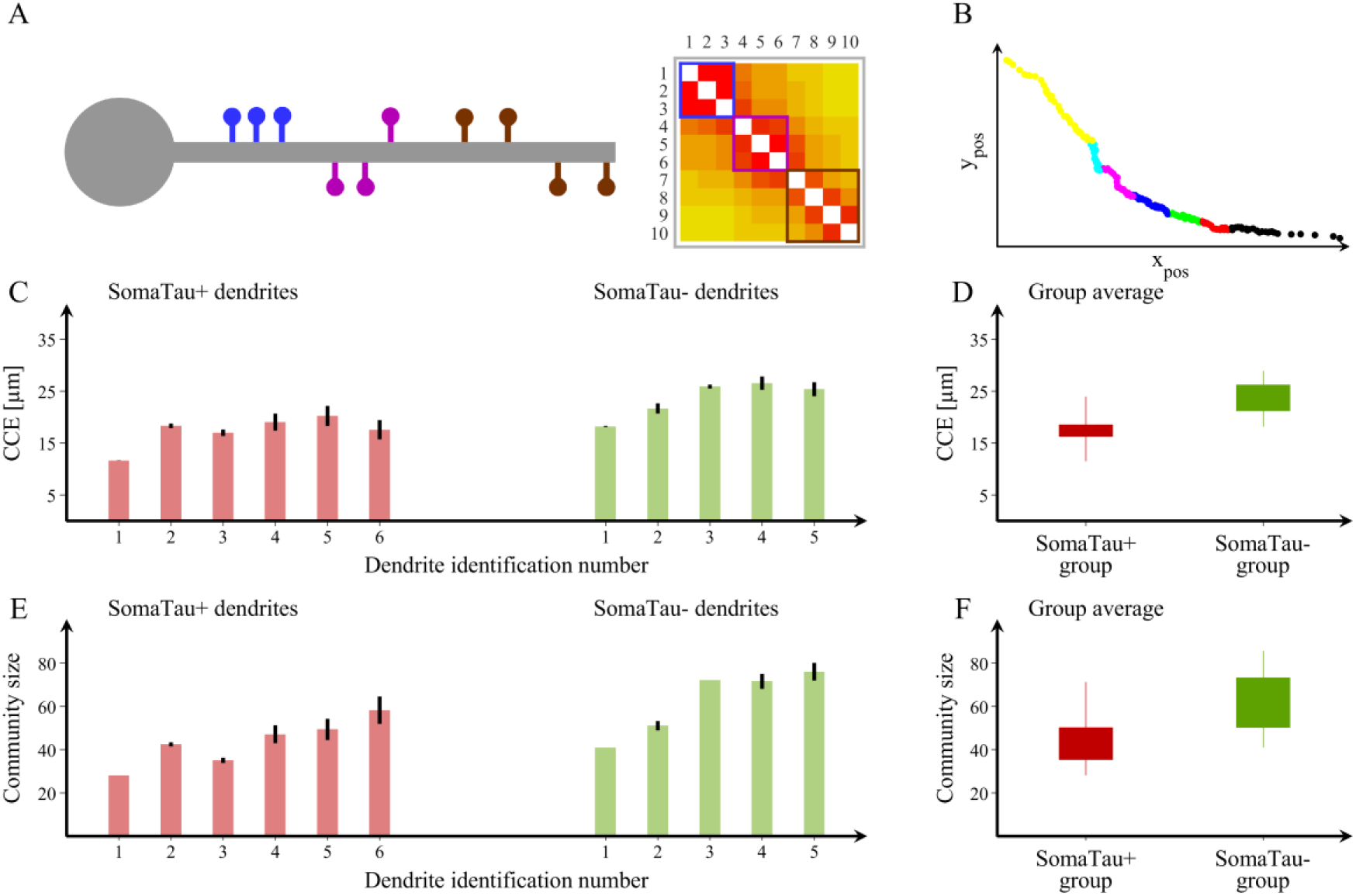
Spine organization into communities. (**A**) Schematic representation of a dendrite with spines that can be clearly separated into three distinct communities and their corresponding adjacency matrix showing the blocks of connections that correspond to each community. (**B**) We also show an example of a community structure for one of the dendrites assessed in the current study with seven communities. The community structure is assessed using the average distance between spines that belong to each community (characteristic community extension, CCE) and the number of spines in each community. Y_pos_ and X_pos_, (μm). (**C, E**) We show the values obtained in these two measures for each SomaTau+ (red, n= 6) and SomaTau- (green, n = 5) dendrite, which are computed by calculating the community structure over 100 trials. In addition, we include (**D**) boxplots with the group averages for the CCE and (**F**) the average community size or number of spines for each community, which are both smaller in SomaTau+ compared to SomaTau- dendrites. The bottom and the top edges of the boxplots denote the 25^th^ and 75^th^ percentiles of the data, respectively. The whiskers extend to the largest and smallest data points. The results are similar after excluding the outlier.

To further investigate this distinctive pattern of organization characterized by smaller and more tightly packed communities of spines in SomaTau+ dendrites compared to SomaTau- dendrites, we conducted two additional analyses. In the first analysis, we assessed whether this organization can be explained by the loss of spines associated with tau pathology, a phenomenon that we have already reported in AD [13]. This analysis showed that the SomaTau+ dendrites have a lower number of spines compared to their SomaTau- counterparts, although these differences did not reach statistical significance (SomaTau+ = 323.0 ± 66.6, SomaTau- = 453.2 ± 135.2, p = 0.11). In the second analysis, we assessed whether SomaTau+ dendrites have a different number of groups of spines that are closely connected with each other, which we refer to as mean grouping coefficient. This analysis showed that the mean grouping coefficient (GC) in SomaTau+ dendrites was significantly higher (*GC*_SomaTau+_ = 0.014 ± 0.003 μm^−1^, *GC*_SomaTau-_= 0.011 ± 0.002 μm^−1^, p < 0.001 (Fig. 4A, 4B), indicating that their spines were organized in more isolated groups along the dendrite, whereas the SomaTau- dendrites have spines that are more evenly spread along the dendrite.

**Fig 4.**
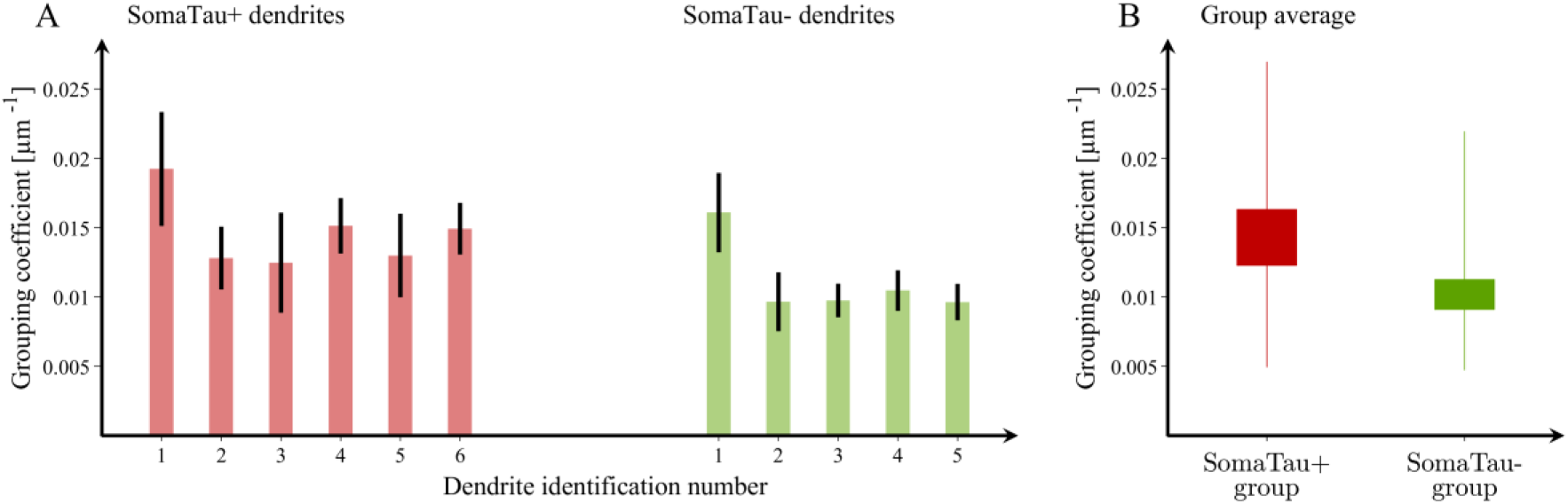
Mean grouping coefficient in dendrites with and without tau pathology. (**A**) Mean grouping coefficient in each single SomaTau+ (n = 6; red) and SomaTau-(n = 5; green) dendrite. Boxplots with the mean grouping coefficients in the SomaTau- and SomaTau+ dendrites. (**B**) The permutation analyses show a higher mean grouping coefficient in the Tau+ compared to the Tau-group (p < 0.001). In all boxplots, their bottom and the top edges denote the 25^th^ and 75^th^ percentiles of the data, respectively. The whiskers extend to the largest and smallest data points. The results are similar after excluding the outliers.

### Replication in two independent AD cases

To assess the generality of these results, we repeated the analyses in two independent AD cases, whose samples were obtained using the same procedures described in the Methods. The first case (P9; gender: male; age: 82 years; cause of death: bronchopneumonia plus heart failure) had 5 SomaTau+ and 2 SomaTau- dendrites with 786 spines, whereas the second case (P14; gender: female; age: 87 years; cause of death: respiratory infection) had 7 SomaTau+ and 3 SomaTau- dendrites with 1376 spines (Supplementary Table S2 and Supplementary Table S3). The results showed that the SomaTau+ dendrites had significantly smaller communities in both the first (*N*_SomaTau+_ = 30.31 ± 9.86 spines, *N*_SomaTau-_= 48.98 ± 32.78 spines, p < 0.001) and second (*N*_SomaTau+_ = 64.32 ± 8.84 spines, *N*_SomaTau-_= 75.85 ± 17.15 spines, p < 0.001) cases. In addition, we also found smaller characteristic community extension in the first (*CCE*_SomaTau+_ = 20.67 ± 7.60 μm, *CCE*_SomaTau-_= 31.10 ± 10.54 μm, p < 0.001) and second (*CCE*_SomaTau+_ = 21.98 ± 3.17 μm, *CCE*_SomaTau-_= 25.65 ± 4.95 μm, p < 0.001) independent cases (Supplementary Fig. S1 and Supplementary Fig. S2). Finally, we found that the spines in SomaTau+ dendrites were part of tighter neighborhoods as illustrated by their higher mean grouping coefficient in both cases (case 1: *GC*_SomaTau+_ = 0.011 ± 0.006 μm^−1^, *GC*_SomaTau-_= 0.007 ± 0.001 μm^−1^, p < 0.001; case 2: *GC*_SomaTau+_ = 0.013 ± 0.003 μm^−1^, *GC*_SomaTau-_= 0.011 ± 0.003 μm^−1^, p < 0.001) (Supplementary Fig. S3 and Supplementary Fig. S4).

Altogether, these results show that the spines of SomaTau+ dendrites are organized into smaller and shorter communities in independent cases, supporting the assumption that spine degeneration is not a random process but follows a specific pattern in response to tau pathology.

### The organization of SomaTau- dendrites becomes similar to SomaTau+ dendrites when clusters of spines are removed

To investigate the mechanisms underlying the more clearly outlined communities and higher number of spine groups in the SomaTau+ dendrites compared to SomaTau- dendrites, we performed a series of simulated attacks on the SomaTau- dendrites from all 3 individuals by progressively removing their spines. We considered three possible attack strategies and assessed their effects on the characteristic community extension as well as on the mean grouping coefficient. These attack strategies are summarized in Fig. 5A, Fig. 5D and Fig. 5G, as well as Fig. 6A, Fig. 6D and Fig. 6G. First, we performed a random attack, by removing multiple single spines at random locations along the dendrite (Fig. 5A, 6A). Second, we performed attacks of blocks of 3 spines, where we removed groups of 3 adjacent spines at random locations along the dendrite (i.e., we select random spines for removal and we remove them together with their first-degree neighbors, Fig. 5D, 6D). Finally, we performed attacks of blocks of 5 spines, where we removed groups of 5 adjacent spines (i.e., we remove randomly selected spines together with their first- and second-degree neighbors, Fig. 5G, 6G). Due to the element of randomness in these attacks, we averaged our results over 100 independent attacks on each dendrite to obtain statistically reliable results.

**Fig 5.**
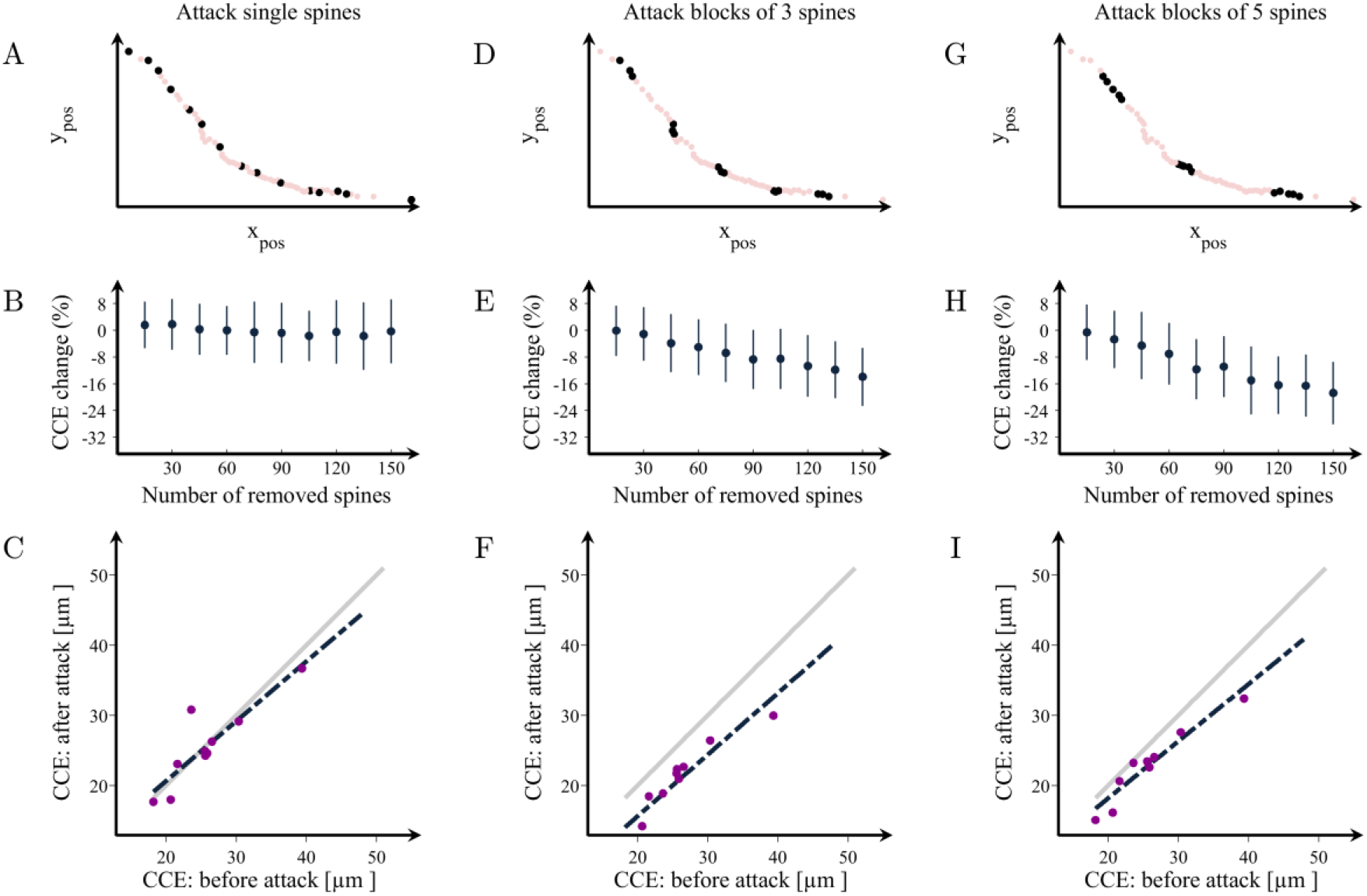
Changes in the organization of dendritic spines after attacks. Examples of attacks on spines in a representative dendrite at (**A**) random locations or in clusters of (**D**) 3 and (**G**) 5 spines. (**B, E, H**) Percentage change in the characteristic community extension (CCE) as a function of the number of removed spines in the three cases, respectively. CCE in the attacked vs. the non-attacked dendrites after the removal of 150 spines in groups of (**C**) 1, (**F**) 3 and (**I**) 5. The grey line shows the theoretical line of no change in the CCE, while the black line is the line that best fits the observed attack data. The values obtained after each attack can also be found in Supplementary Table S4. Means and standard deviations are computed over 100 random attacks.

**Fig 6.**
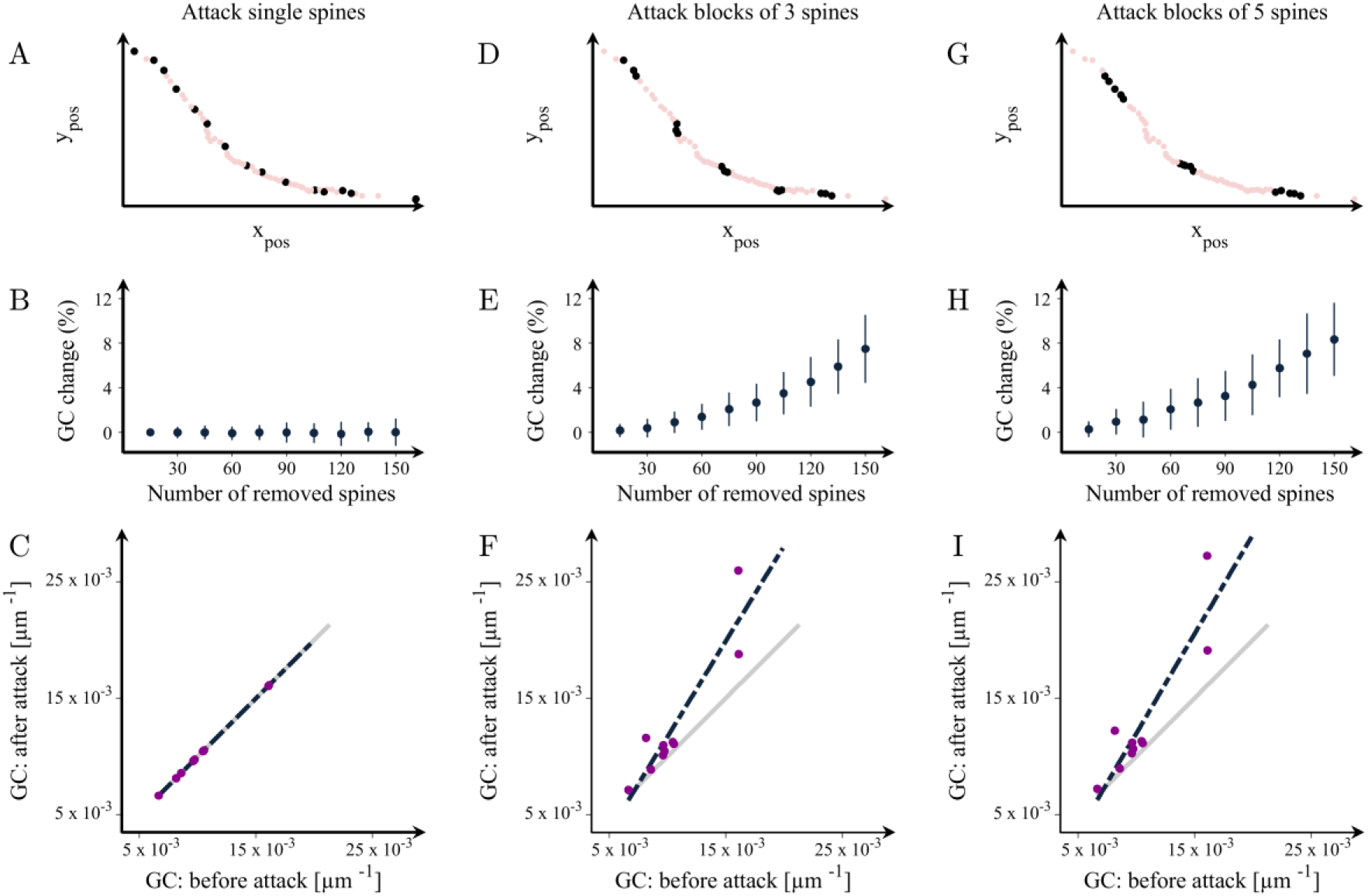
Changes in the mean grouping coefficient after random attacks of spines in healthy dendrites. Illustration of attacks on random spines in a representative dendrite in groups of (**A**) 1, (**D**) 3 and (**G**) 5. (**B, E, H**) Percentage increases in the mean grouping coefficient (GC) in that dendrite as a function of the number of removed spines in the three cases respectively. Mean grouping coefficient of the attacked vs. healthy dendrites after the removal of 150 spines in groups of (**C**) 1, (**F**) 3 and (**I**) 5. The grey line shows the theoretical line of no change in the grouping coefficient, while the black line is the line that best fits the observed data after the attacks. This figure shows results for dendrite no. 2; the complete set of results for each SomaTau-dendrite are shown in Supplementary Table S5. Means and standard deviations are computed over 100 random attacks.

These analyses show that removing random single spines from the dendrite had no effect on the communities and groups of spines, even when a large number of spines was removed (Fig. 5B, 5C, 6B, 6C). In contrast, when the spines are removed in blocks of 3 or 5, the attacked SomaTau- dendrites show a smaller characteristic community extension (Fig. 5E-5I, Supplementary Table S4) and a greater number of groups denoted by increases in the mean grouping coefficient (Fig. 6E-6I, Supplementary Table S5), similarly to the SomaTau+ dendrites. These analyses indicate that the mechanism underlying the distinct organization pattern in dendrites with tau pathology entails the loss of blocks of spines.

## DISCUSSION

The understanding of how spines are lost in AD is still unclear but could potentially reveal the mechanisms underlying synaptic damage in this disorder. In this study, we show that the presence of tau pathology in CA1 pyramidal neurons is associated with a distinct organization of spines into smaller and shorter communities. This reorganization is due to a specific pattern of spine loss in groups or blocks of spines in SomaTau+ dendrites. These findings indicate that the loss of spines associated with tau pathology is not a random process but occurs in clusters, shedding light onto the neurodegenerative changes that occur in the course of AD.

It is well established that the tau protein plays an important role in stabilizing microtubules and regulating axonal transport [22]. In addition to its role in supporting microtubules, tau also regulates other processes associated with synaptic function, being detected in the dendrites and the postsynaptic structures of healthy neurons [23–25]. In particular, tau can directly interact with scaffolding proteins, regulating the targeting of glutamatergic receptors to postsynaptic sites in spines. Moreover, there is evidence showing that tau is involved in long-term depression in the CA1 of the hippocampus [26, 27]. Overall, these studies suggest that tau exerts a central role controlling the normal healthy function of the synapses in the brain.

In AD, tau undergoes pathological changes such as hyperphosphorylation, which affect its affinity towards microtubules [28]. These changes eventually lead to the detachment of tau from microtubules, their translocation from axons to the somatodendritic compartment and spines, where it can interfere with synaptic function [29]. The modified tau molecules tend to self-assemble into paired helical filaments, which form neurofibrillary tangles that exert toxicity on the neuron [29], leading to neurodegenerative changes such as the loss of spines. In line with this, we have previously found a loss of spines in pyramidal neurons with tau pathology in AD individuals [13]. Since generally one spine corresponds to one excitatory synapse [30], the loss of spines in pyramidal neurons with tau indicates they receive a lower number of excitatory inputs, reducing synaptic communication.

However, to this date, the mechanisms underlying spine loss in AD have not been investigated. In line with this, there is increasing evidence showing that functionally related synapses are organized in clusters and that these clusters are crucial for cognitive functions that become impaired in AD such as memory storage and maintenance [11, 12]. This has led to the proposal of a hypothesis based on clustered plasticity, which supports the idea that the number and position of spines in the dendrites as well as their excitatory synapses depend on the dynamics of neuronal connectivity and that learning and memory can lead to the organization of spines in clusters [31, 32]. Thus, in this study we assessed the spine organization in a unique dataset of dendrites from neurons with and without tau pathology from one AD case. By modeling dendrites as graphs, where the spines represent nodes and the connections between the spines reflect how close they are to each other, we have found a specific change in the pattern of spine organization into shorter and smaller communities in dendrites with tau pathology. To assess whether these findings were generalizable to other AD cases, we repeated the analyses in two independent individuals and found the same results, suggesting that the observed changes in spine organization are consistent across patients.

To investigate the mechanisms responsible for this abnormal organization, we performed a series of simulated attacks to the dendrites without tau pathology. These attacks revealed that only by removing spines in clusters, the attacked SomaTau- dendrites show an organization similar to that of SomaTau+ dendrites. These findings are in line with previous studies showing that the organization of spines in clusters is biologically meaningful in healthy neurons [9–12]. Here, we extend these findings by confirming that this organization is also associated with pathological conditions such as AD. In particular, our results show that AD is associated with a loss of clusters of spines, which could be the mechanism by which tau drives synaptic damage in this disorder, leading the way to cognitive deficits. With increasing recognition of AD as a synaptic disorder [33], maintaining the function of spines may become an important therapeutic target in the future. Our results suggest that when it comes to the normal organization of the dendrite, it is important to maintain the healthy function of clusters of spines rather than single spines, and thus new treatments should focus on preventing damage to these relevant functional units. In addition, recent clinical trials in sporadic and familial AD show that anti-amyloid drugs reduce the levels of phosphorylated tau, indicating that these treatments have downstream effects on tau metabolism and synaptic function [34]. Thus, assessing the organization of spines or developing in-vivo biomarkers that reflect the integrity of spine clusters within the dendrites will become important to assess the clinical effects of these trials since synapses are crucial for memory and other cognitive functions that become impaired in AD.

Our approach of modeling the dendrites as a graph has several advantages. By integrating each spine in a network with all the other spines of the same dendrite, we can assess its architecture using topological measures that reflect the communities. These measures have been extensively applied to assess the organization of the brain in human individuals, revealing that normal brain function requires a network organization divided into communities, with potentially different functions or connectivity patterns [35, 36]. To our knowledge, our study is the first to apply a graph-theory approach to assess the organization of spines. This approach allowed us to establish that the random loss of spines is not responsible for the reorganization observed in dendrites with tau pathology. Instead, clusters of spines seem to play a key role in this reorganization, in line with previous studies using magnetic resonance imaging showing that groups of tightly connected brain regions are an important measure that becomes altered in AD individuals [37].

Our study has also a few limitations such as the lack of ante-mortem cognitive measures for the AD cases that we could have used to relate to the loss of spine blocks and assess the clinical value of these changes. In addition, although the number of dendrites examined in the study may appear low, one should take into account that brain tissue for intracellular injections has to be obtained with short postmortem delays (less than 5 h), with the total number of cells that can be injected being relatively few since it takes approximately 10 minutes to inject a cell and the best injections are obtained within a relatively short time window of 24–48 h after fixation. The lack of brain tissue from elderly individuals without AD would have been interesting to include in order to assess the organization of spines in normal brains, although normal brains are likely to have several other pathologies that could influence synaptic integrity [38]. However, our approach of comparing neighboring cells with and without neurofibrillary pathology within each individual is the best approach to avoid confounding factors such as: (1) morphological differences in the structure of pyramidal cells due to regional specializations (i.e., pyramidal cells in different cortical regions and layers may show morphological differences); (2) high inter-individual variability (sex, age, medical treatment, etc.), factors that could affect brain structure; and (3) the highly variable course of AD, as the neuropathological changes are not homogenous among AD individuals or in different regions of the brain of the same individual, giving rise to variation in the alterations to cortical circuits.

In summary, this study shows that AD is associated with a reorganization of spines into smaller and more tightly packed communities due to a loss of groups of spines in dendrites with tau pathology. These findings suggest that tau targets spines in clusters along the dendrite, damaging synaptic connections that potentially share the same synaptic contact. Future studies should validate our results in larger numbers of individuals and in additional brain regions.

## METHODS

### Human brain samples

Brain tissue was obtained at autopsy from the Instituto de Neuropatología (Dr I. Ferrer, Servicio de Anatomía Patológica, IDIBELL-Hospital Universitario de Bellvitge, Barcelona, Spain), from the Banco de Tejidos Fundación CIEN (Centro Alzheimer, Fundación Reina Sofía, Madrid, Spain) and from the Neurological Tissue Bank (Biobanc-Hospital Clínic-IDIBAPS, Universidad de Barcelona, Spain)(Supplementary Table S2). Following neuropathological examination, the pathological state of the AD individual (P13; gender: male; age: 83 years; cause of death: respiratory failure) was defined as AD VI/C and LB according to the CERAD (*The Consortium to Establish a Registry for Alzheimer’s Disease (CERAD) | Neurology*, n.d.) and the Braak and Braak criteria [39]. The time between death and tissue fixation was maximum 3 h.

The brain tissue was obtained following national laws and international ethical and technical guidelines on the use of human samples for biomedical research purposes. In all cases, brain tissue donation, processing and use for research were performed in compliance with published protocols [40], which include the obtaining of informed consent for brain tissue donation from living donors, and the approval of the whole donation process by local ethical committees (Comité de Bioseguridad IDIBELL, Comité de Ética de la Investigación del Instituto de Salud Carlos III and Comité de Ética BioBanc, respectively).

### Intracellular injections and immunocytochemistry

Coronal sections (250 μm) were obtained with a vibratome and labeled with 4,6 diamino-2-phenylindole (DAPI; Sigma, St Louis, MO, USA) to identify cell bodies. Pyramidal cells from the CA1 region were then individually injected with Lucifer Yellow (LY) (8% in 0.1 M Tris buffer, pH 7.4). LY was applied to each injected cell by continuous current until the distal tips of each cell fluoresced brightly, indicating that the dendrites were completely filled and ensuring that the fluorescence did not diminish at a distance from the soma. Following intracellular injections, all sections were immunostained with both, anti-LY,anti- phospho-tau_AT8_ and anti-phospho-tau_PHF-1_ antibodies [13]. We performed a total of 62 intracellular injections in CA1 pyramidal neurons, and after excluding the neurons that were not appropriated for the analysis (see Introduction), 6 were used for 3D reconstruction. We reconstructed the spines in three dimensions by confocal microscopy using a previously described methodology [41]. Only complete dendrites were included in the analysis. It should be noted that the intracellular injections have to be performed before the immunostaining with anti- phospho-tau antibodies. Therefore, it is not possible to know which neurons present neurofibrillar pathology before the intracellular injections.

### Construction of graphs

We modeled each single dendrite as a graph represented by a set of nodes and edges, where nodes represent the elements of the graph (dendritic spines) and the edges indicate the strength of association between these elements. In this model, the strength of association between the nodes is proportional to the physical closeness between the spines and is calculated as the inverse distance between the spines along the dendrite. Let *i* and *k* represent two spines along the dendrite, and let *S* = *i* → *j* → *a* → *b* → ⋯ → *u* → *v* → *k* denote the complete sequence of neighboring spines positioned between *i* and*k*. For each pair of neighboring spines belonging to this sequence, e.g. *u* and *v* at positions *r*_*u*_ = (*x*_*u*_, *y*_*u*_, *z*_*u*_) and *r*_*v*_ = (*x*_*v*_, *y*_*v*_, *z*_*v*_) respectively, the distance between them is calculated as:

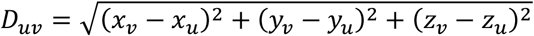

Since the distance between any pair of neighboring spines along the sequence *S* is much smaller than the total length of the dendrite (*D*_*uv*_ ≪ *L*), the total distance between *i* and *k* along the dendrite is represented as a sum of the distances between all consecutive pairs of spines within the sequence:

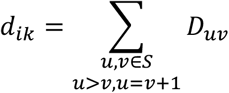

Finally, the strength of association between spines *i* and *k*can be expressed as

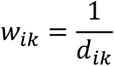

After calculating the pairwise strength of association between all possible pairs of spines and the construction of the corresponding graph, various graph measures can be calculated in order to investigate its local and global organization. In our analysis, we focused on the global community organization of the dendrite by calculating the number of spines within each community as well as the characteristic community extension. In addition, we also calculated the total number of spines of each dendrite and the mean grouping coefficient.

### Community structure and characteristic community extension

The community structure of a graph reflects how well the graph can be fragmented into different sub-graphs or communities. Communities are defined as groups of nodes that are tightly connected with each other but are poorly connected with nodes from other communities. Here, we calculate the community structure by using the Louvain algorithm [21], which optimizes the modularity of the graph by iteratively merging communities into single nodes and subsequently recalculating the modularity on the corresponding graph. Modularity is a measure that compares the density of within-community connections with that of a random graph; higher modularity values indicate a better division of the graph into communities. The modularity is calculated as:

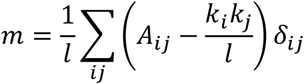

where *l* is the number of edges in the graph, *A*_*ij*_ is the adjacency matrix of the graph, *k*_*i*_ (*k*_*j*_) is the degree of the node *i* (*j*), *δ*_*ij*_ is 1 if the two nodes belong to the same community and 0 otherwise, and the sum is performed over all pairs of nodes in the graph. Due to the inherent randomness of the Louvain algorithm, the community structure measures are calculated as averages of 100 independent runs of the algorithm.

The community structure of the dendrites can be characterized by the average size and the characteristic extension of the communities present within a dendrite. The community size is defined as the number of spines present within the same community. Furthermore, we calculate the characteristic community extension (in *μm* as the average distance between all pairs of spines that belong to the same community:

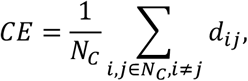

where *i* and *j* are two spines at a distance *d*_*ij*_ that belong to the same community *C*,and *N*_*C*_ is the total number of spines in the community *C*. Finally, the characteristic community extension (CCE) for a given dendrite is defined as the average community extensions of all communities within the corresponding dendrite.

### Mean grouping coefficient (GC)

For each spine, we quantified the degree to which that node is part of a tightly connected neighborhood. Given a set of three spines *i*, *j* and *k*, let the connection strength between nodes *i* and*j* be represented as *w*_*ij*_, between nodes *i* and *k* as *w*_*ik*_, and between nodes *j* and *k* as *w*_*jk*_. The weight of the triangle formed by these three connections can be expressed as:

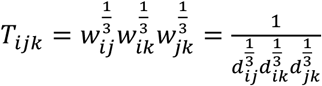

where *d*_*ij*_, *d*_*ik*_ and *d*_*jk*_ are the distances between the corresponding spines. The triangle weight *T*_*ijk*_ reflects the positions of the three spines along the dendrite relative to each other. Therefore, small values of *T*_*ijk*_ indicate that the three corresponding spines *i*, *j* and *k* are spaced far away from each other, while larger values indicate that the three spines are positioned closely to each other along the dendrite.

The mean grouping coefficient for the spine *i* is the average weight of all triangles the spine *i* is part of, and it characterizes the position of spine *i* in the dendrite relative to all other spines. This coefficient ranges between 0 and 1, where 1 is calculated for spines that are part of a tight neighborhood with many spines positioned nearby. Conversely, spines with a coefficient of 0 belong to a spread-out neighborhood and are as far as possible from the other spines. We calculated this coefficient as:

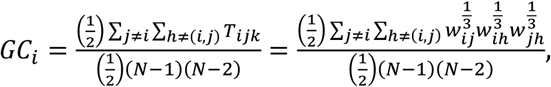

where *N* is the number of spines along the dendrite to which *i* belongs to.

### Statistical comparison of the results

Differences between SomaTau+ and SomaTau- dendrites were assessed using non-parametric permutation tests with 10 000 permutations, which were considered significant for a two-tailed test of the null hypothesis at p<0.05 [42]. The tests were performed by first calculating the difference in the means between the two groups. Then, we randomly permuted the elements from both groups and calculated the differences in the means between the new randomized groups. By repeating this procedure multiple times, we obtained a null distribution of between-group differences. Finally, we obtained the two-tailed p-value as the proportion of between-group differences in the null distribution that are greater than the absolute value of the original difference. The community structure parameters were obtained as averages of 100 trials.

## Acknowledgements

This work was supported by grants from the Spanish Ministerio de Ciencia, Innovación y Universidades (grant IJCI-2016-27658 to PMS, grant PGC2018-094307-B-I00), the Cajal Blue Brain Project (the Spanish partner of the Blue Brain Project initiative from EPFL, Switzerland) and the Centro de Investigación Biomédica en Red sobre Enfermedades Neurodegenerativas (CIBERNED, CB06/05/0066, Spain). JBP is currently supported by grants from the Swedish Research Council (#2018-02201), Hjärnfonden (#FO2019-0289), Alzheimerfonden (#AF-930827) and the Strategic Research Programme in Neuroscience at Karolinska Institutet (Stratneuro Startup Grant). We would like to thank Carmen Álvarez and Lorena Valdés for their helpful assistance.

## Author contributions

P.M.S., J.D., I.F., J.B.P. and G.V. conceptualized the study. P.M.S., J.D., I.F. performed the intracellular injections and immunocytochemistry analyses. M.M. performed the graph and statistical analyses. All authors contributed to drafting, reviewing and editing the manuscript.

## Supplementary Information

SI_Appendix: This appendix contains Supplementary Tables S1-S5 and Supplementary Figures S1-S4.

## Data availability

All relevant data is made available within the manuscript and the associated Supplementary Information files.

