## Supplementary Information for "Dendritic spines are lost in clusters in Alzheimer’s disease"

**Supplementary Table S1: Summary of the morphological properties of all SomaTau+ and SomaTau- dendrites used in the analysis for individual P13.**
* Indicates the presence of phospho-tau in the distal segment of the dendrite.

| | Dendrite identification number | Cell identification number | Individual | Number of spines | Dendritic length (μm) | Spine density |
|---|---|---|---|---|---|---|
| Soma Tau+ | **1** | 37 | P13 | 225 | 166,63 | 1,35 |
| | **2** | 37 | P13 | 296 | 228,15 | 1,29 |
| | **3** | 37 | P13 | 319 | 211,41 | 1,51 |
| | **4*** | 49 | P13 | 319 | 186,54 | 1,71 |
| | **5** | 74 | P13 | 351 | 250,17 | 1,4 |
| | **6** | 43 | P13 | 428 | 191,7 | 2,24 |
| Soma Tau- | **1** | 81 | P13 | 246 | 172,85 | 1,43 |
| | **2** | 25 | P13 | 403 | 224,71 | 1,79 |
| | **3** | 25 | P13 | 505 | 234,36 | 2,15 |
| | **4** | 25 | P13 | 512 | 245,74 | 2,08 |
| | **5** | 25 | P13 | 600 | 242,13 | 2,47 |

**Supplementary Table S2. Summary of the AD individual data and the LY-injected neurons in CA1 region.** In brackets are indicated the number of neurons used for 3D examination. Aβ, Amyloid-β; LB, Lewy bodies; NF, Neurofibrillar.

| **AD Individual** | **P9** | **P13** | **P14** |
|---|---|---|---|
| **Age/Gender** | Male/82 | Male/83 | Female/87 |
| **NF/Aβ pathology Braak stage** | AD V/C | AD VI/C and LB | AD III-IV /0-A |
| **Post-mortem delay (h)** | 3 | 2:30 | 1:30 |
| **Cause of death** | Bronchopneumonia plus heart failure | Respiratory failure | Respiratory infection |
| **CA1 LY-injected pyramidal neurons** | 40 (15) | 62 (8) | 48 (8) |

**Supplementary Table S3: Summary of the properties of all SomaTau+ and SomaTau- dendrites used in the analysis for individuals P9 and P14.** * Indicates the presence of phospho-tau in the distal segment of the dendrite

| | Dendrite identification number | Cell identification number | Individual | Number of spines | Dendritic length (µm) | Spine density |
|---|---|---|---|---|---|---|
| Soma Tau+ | **1*** | 197 | P9 | 179 | 177,38 | 1,01 |
| | **2*** | 197 | P9 | 203 | 196,64 | 1,03 |
| | **3** | 2 | P9 | 270 | 172,02 | 1,56 |
| | **4** | 195 | P9 | 299 | 225,65 | 1,32 |
| | **5** | 195 | P9 | 355 | 222,4 | 1,59 |
| Soma Tau- | **1** | 181 | P9 | 258 | 206,3 | 1,25 |
| | **2** | 198 | P9 | 475 | 253,84 | 1,87 |
| Soma Tau+ | **1** | 239 | P14 | 337 | 187,28 | 1,8 |
| | **2*** | 173 | P14 | 402 | 250,36 | 1,6 |
| | **3*** | 173 | P14 | 411 | 233,31 | 1,76 |
| | **4*** | 121 | P14 | 432 | 246,6 | 1,75 |
| | **5** | 5 | P14 | 447 | 177,1 | 2,52 |
| | **6** | 121 | P14 | 450 | 249,29 | 1,08 |
| | **7** | 132 | P14 | 623 | 209,15 | 2,98 |
| Soma Tau- | **1** | 13 | P14 | 363 | 208,97 | 1,74 |
| | **2** | 258 | P14 | 573 | 208,43 | 2,75 |
| | **3** | 141 | P14 | 693 | 221,68 | 3,13 |

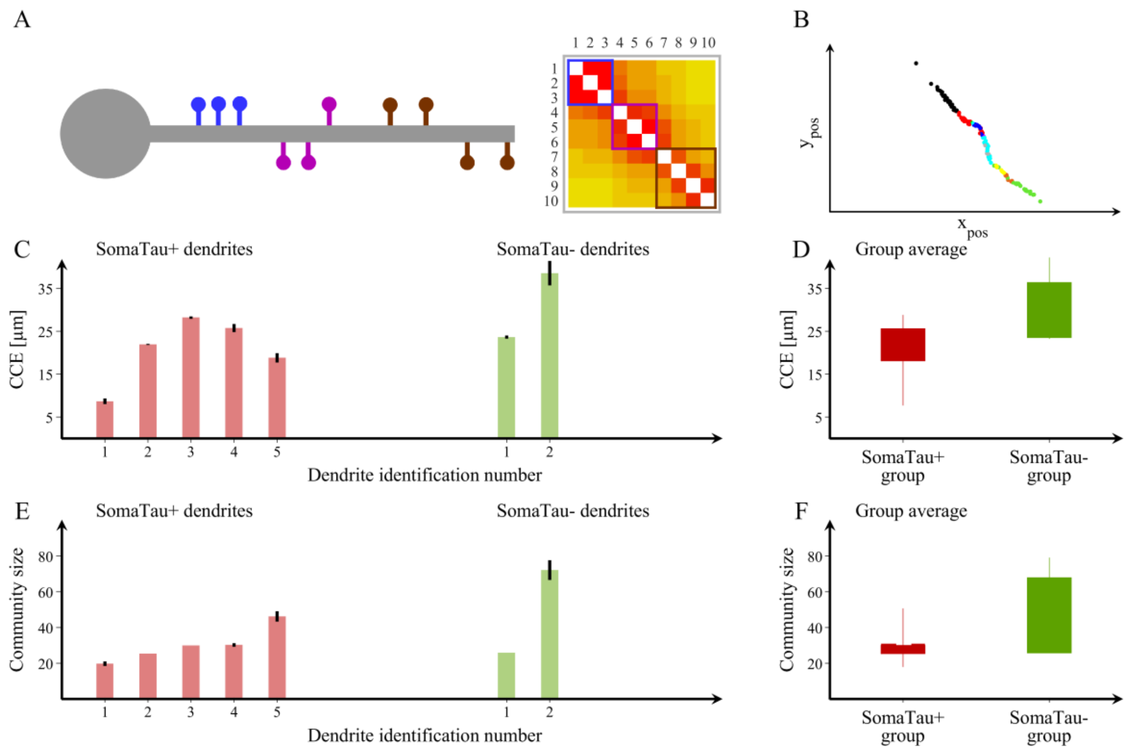

**Supplementary Figure S1. Spine organization into communities for individual P9.**

(**A**) Schematic representation of a dendrite with spines that can be clearly separated into three distinct communities and their corresponding adjacency matrix showing the blocks of connections that correspond to each community. (**B**) We also show an example of a community structure for one of the dendrites assessed in the current study with seven communities. The community structure is assessed using the average distance between spines that belong to each community (characteristic community extension, CCE) and the number of spines in each community. $Y_{pos}$ and $X_{pos}$, (µm). (**C, E**) We show the values obtained in these two measures for each SomaTau+ (red, n= 5) and SomaTau- (green, n = 2) dendrite, which are computed by calculating the community structure over 100 trials. In addition, we include (**D**) boxplots with the group averages for the CCE and (**F**) the average community size or number of spines for each community, which are both smaller in SomaTau+ compared to SomaTau- dendrites. The bottom and the top edges of the boxplots denote the 25$^{th}$ and 75$^{th}$ percentiles of the data, respectively. The whiskers extend to the largest and smallest data points. The results are similar after excluding the outlier.

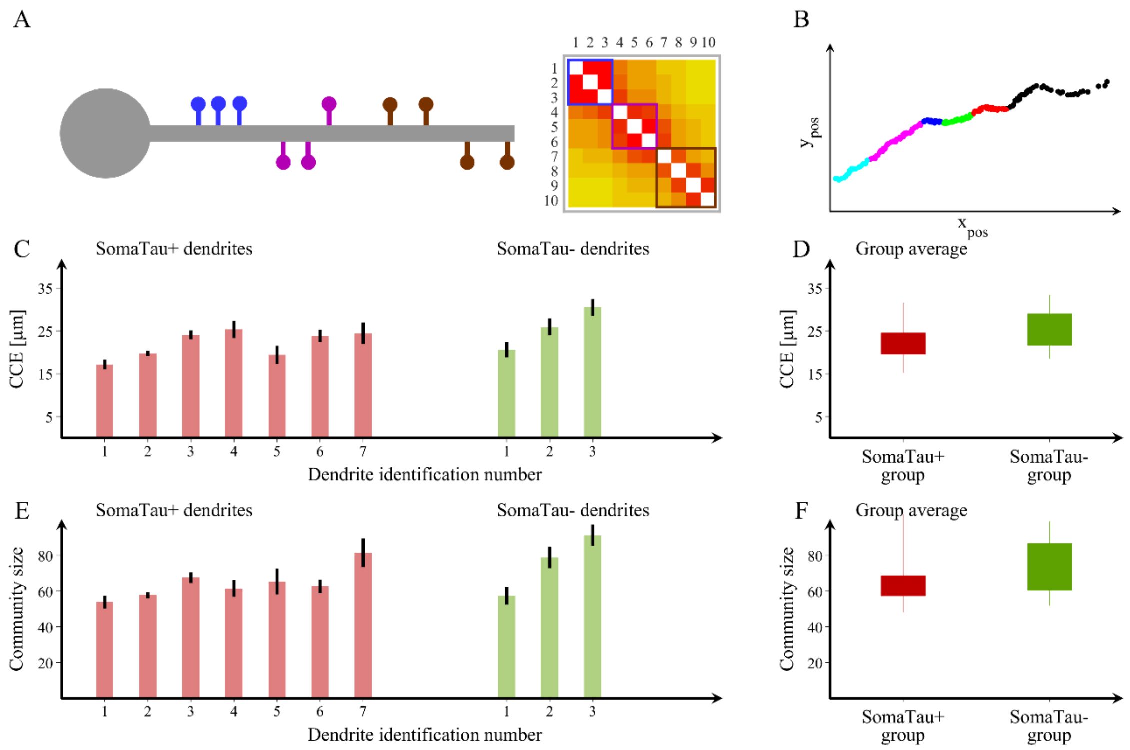

**Supplementary Figure S2. Spine organization into communities for individual P14.**

(**A**) Schematic representation of a dendrite with spines that can be clearly separated into three distinct communities and their corresponding adjacency matrix showing the blocks of connections that correspond to each community. (**B**) We also show an example of a community structure for one of the dendrites assessed in the current study with seven communities. The community structure is assessed using the average distance between spines that belong to each community (characteristic community extension, CCE) and the number of spines in each community $Y_{pos}$ and $X_{pos}$, (μm). (**C, E**) We show the values obtained in these two measures for each SomaTau+ (red, n= 7) and SomaTau- (green, n = 3) dendrite, which are computed by calculating the community structure over 100 trials. In addition, we include (**D**) boxplots with the group averages for the CCE and (**F**) the average community size or number of spines for each community, which are both smaller in SomaTau+ compared to SomaTau- dendrites. The bottom and the top edges of the boxplots denote the 25$^{th}$ and 75$^{th}$ percentiles of the data, respectively. The whiskers extend to the largest and smallest data points. The results are similar after excluding the outlier.

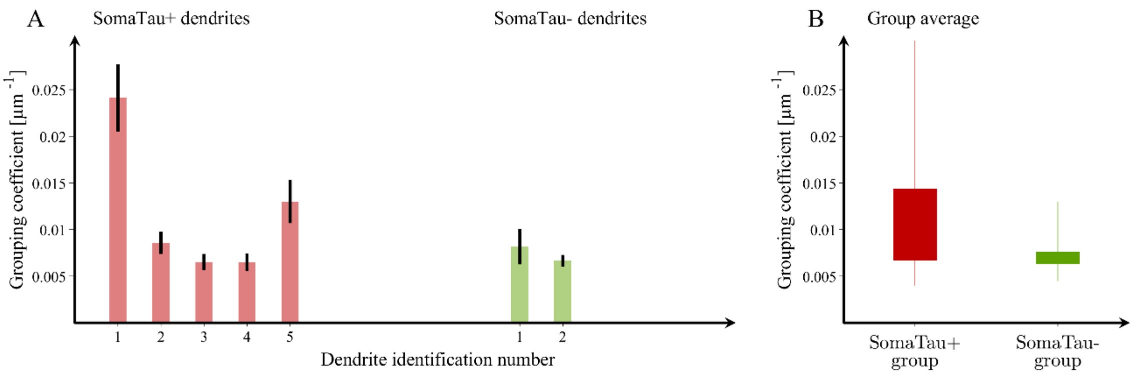

**Supplementary Figure S3. Mean grouping coefficient in dendrites with and without tau pathology for individual P9.**

(**A**) Mean grouping coefficient in each single SomaTau+ (n = 5; red) and SomaTau- (n = 2; green) dendrite. Boxplots with the mean grouping coefficients in the SomaTau- and SomaTau+ dendrites. (**B**) The permutation analyses show a higher mean grouping coefficient in the Tau+ compared to the Tau- group ($p < 0.001$). In all boxplots, their bottom and the top edges denote the 25$^{th}$ and 75$^{th}$ percentiles of the data, respectively. The whiskers extend to the largest and smallest data points. The results are similar after excluding the outliers.

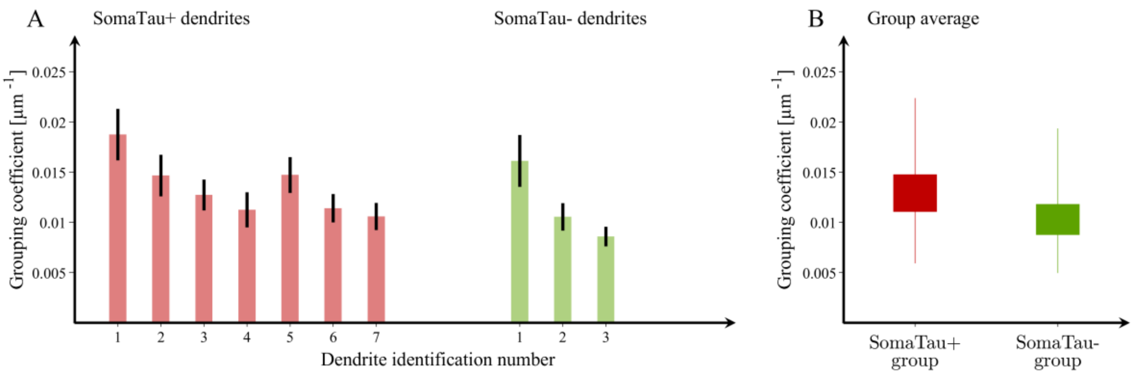

**Supplementary Figure S4. Mean grouping coefficient in dendrites with and without tau pathology for individual P14.**

(**A**) Mean grouping coefficient in each single SomaTau+ (n = 7; red) and SomaTau- (n = 3; green) dendrite. Boxplots with the mean grouping coefficients in the SomaTau- and SomaTau+ dendrites. (**B**) The permutation analyses show a higher mean grouping coefficient in the Tau+ compared to the Tau- group ($p < 0.001$). In all boxplots, their bottom and the top edges denote the 25th and 75th percentiles of the data, respectively. The whiskers extend to the largest and smallest data points. The results are similar after excluding the outliers.

**Supplementary Table S4: Percentage change in the characteristic community extension (CCE) for all SomaTau- dendrites as a function of the number of removed spines.** The changes are shown in the cases when multiple spines are removed from the dendrites individually, in blocks of 3 and in blocks of 5 spines. The means are followed by standard deviations calculated over 100 trials.

| **Attack by removal of individual spines** | | | | | |
|---|---|---|---|---|---|
| **Number of removed spines** | **15** | **30** | **45** | **60** | **75** |
| **Dendrite 1 (P13)** | 0.0(2.9) | -1.5(5.2) | -1.0(8.4) | -2.9(7.8) | -3.1(9.1) |
| **Dendrite 2 (P13)** | 5.9(9.4) | 6.0(10.9) | 7.4(9.6) | 7.8(9.3) | 10.0(10.9) |
| **Dendrite 3 (P13)** | -0.5(3.8) | -0.7(5.8) | -1.1(7.4) | -2.0(7.9) | -2.3(7.5) |
| **Dendrite 4 (P13)** | 1.6(6.9) | 1.8(7.6) | 0.3(7.7) | 0.0(7.3) | -0.6(9.1) |
| **Dendrite 5 (P13)** | 0.1(8.6) | -1.1(8.2) | -0.4(7.8) | 0.4(8.3) | -1.0(8.8) |
| **Dendrite 1 (P9)** | 5.4(9.0) | 9.6(11.2) | 7.2(12.1) | 9.4(11.6) | 11.1(12.2) |
| **Dendrite 2 (P9)** | 0.0(11.2) | -1.9(12.1) | -4.1(11.6) | -3.6(12.2) | -4.8(12.3) |
| **Dendrite 1 (P14)** | -2.2(13.1) | -5.4(11.7) | -5.3(11.9) | -5.9(11.7) | -6.9(13.1) |
| **Dendrite 2 (P14)** | 0.5(10.1) | -0.1(10.9) | 0.7(10.0) | -0.1(10.7) | -0.1(11.4) |
| **Dendrite 3 (P14)** | 1.3(10.7) | -0.4(10.3) | 0.3(10.5) | -1.7(9.9) | -2.9(9.8) |
| **Number of removed spines** | **90** | **105** | **120** | **135** | **150** |
| **Dendrite 1 (P13)** | -3.4(11.9) | -4.7(12.5) | -4.4(11.0) | -3.3(12.6) | -0.8(16.9) |
| **Dendrite 2 (P13)** | 6.7(12.8) | 8.2(9.9) | 7.8(12.1) | 7.3(11.6) | 7.7(11.1) |
| **Dendrite 3 (P13)** | -2.0(8.9) | -2.0(9.6) | -0.7(9.5) | -1.9(9.1) | -4.3(9.2) |
| **Dendrite 4 (P13)** | -0.8(9.0) | -1.7(7.6) | -0.5(9.5) | -1.7(10.1) | -0.3(9.5) |
| **Dendrite 5 (P13)** | 0.1(7.9) | -0.7(8.6) | -2.1(8.6) | -1.0(9.2) | -1.8(8.5) |
| **Dendrite 1 (P9)** | 17.5(16.4) | 21.5(16.9) | 22.7(18.5) | 26.4(16.6) | 32.5(20.0) |
| **Dendrite 2 (P9)** | -5.8(11.5) | -5.8(11.4) | -4.6(11.7) | -7.3(11.9) | -4.9(14.8) |
| **Dendrite 1 (P14)** | -6.9(12.8) | -9.1(12.3) | -9.1(12.8) | -10.3(12.1) | -11.3(12.6) |
| **Dendrite 2 (P14)** | -0.3(10.5) | -1.0(10.3) | -2.9(11.3) | -2.9(11.2) | -4.2(11.1) |
| **Dendrite 3 (P14)** | -2.7(11.6) | -3.1(12.3) | -3.2(10.1) | -3.8(9.7) | -2.7(11.5) |
| **Attack by removal of spines in blocks of 3** | | | | | |
| **Number of removed spines** | **15** | **30** | **45** | **60** | **75** |
| **Dendrite 1 (P13)** | -3.0(6.5) | -7.5(7.6) | -8.3(10.1) | -10.7(10.7) | -14.2(11.0) |
| **Dendrite 2 (P13)** | 3.6(8.3) | 4.0(9.5) | 2.8(9.9) | 1.8(10.9) | -0.5(11.3) |
| **Dendrite 3 (P13)** | -2.3(6.9) | -2.7(7.1) | -5.8(9.3) | -8.2(8.0) | -11.0(7.9) |
| **Dendrite 4 (P13)** | -0.1(7.5) | -1.1(8.1) | -3.8(8.7) | -5.0(8.3) | -6.8(8.8) |
| **Dendrite 5 (P13)** | -1.8(9.1) | -1.1(8.1) | -3.0(8.0) | -2.5(9.1) | -5.2(9.8) |
| **Dendrite 1 (P9)** | 3.7(7.6) | 2.8(9.0) | 3.2(12.0) | 1.5(13.8) | 0.7(12.3) |
| **Dendrite 2 (P9)** | -4.9(11.5) | -6.2(12.9) | -10.1(11.1) | -11.5(11.1) | -14.5(9.9) |
| **Dendrite 1 (P14)** | -5.4(11.2) | -7.9(11.1) | -10.3(10.7) | -14.7(10.4) | -16.6(10.7) |
| **Dendrite 2 (P14)** | 0.9(11.5) | -0.9(11.2) | -2.6(10.4) | -5.0(10.6) | -5.2(11.7) |
| **Dendrite 3 (P14)** | -1.9(9.7) | -1.8(10.4) | -3.8(11.0) | -4.7(10.0) | -5.3(11.0) |

| Number of removed spines | 90 | 105 | 120 | 135 | 150 |
|---|---|---|---|---|---|
| **Dendrite 1 (P13)** | -16.0(11.1) | -21.5(10.9) | -25.6(11.1) | -33.5(9.3) | -39.0(12.4) |
| **Dendrite 2 (P13)** | -3.6(10.8) | -7.0(9.7) | -8.5(10.8) | -10.5(10.3) | -13.8(9.1) |
| **Dendrite 3 (P13)** | -12.1(7.8) | -13.3(8.7) | -12.5(8.5) | -15.8(8.8) | -18.4(8.6) |
| **Dendrite 4 (P13)** | -8.7(8.8) | -8.5(9.0) | -10.7(9.3) | -11.8(8.5) | -13.9(8.7) |
| **Dendrite 5 (P13)** | -7.4(9.7) | -8.5(8.5) | -10.6(9.8) | -13.1(8.4) | -13.9(10.0) |
| **Dendrite 1 (P9)** | -0.3(15.2) | -2.8(14.3) | -8.4(14.0) | -13.5(14.2) | -18.9(13.1) |
| **Dendrite 2 (P9)** | -16.5(10.8) | -16.9(10.0) | -19.5(10.6) | -21.7(9.7) | -22.8(9.8) |
| **Dendrite 1 (P14)** | -20.8(9.3) | -23.5(10.4) | -25.8(10.3) | -27.5(10.8) | -30.2(9.3) |
| **Dendrite 2 (P14)** | -7.5(11.4) | -9.4(9.8) | -9.6(11.5) | -11.4(10.7) | -11.8(10.5) |
| **Dendrite 3 (P14)** | -8.0(10.1) | -9.2(10.4) | -9.9(11.2) | -11.6(10.2) | -11.6(11.6) |

**Attack by removal of spines in blocks of 5**

| Number of removed spines | 15 | 30 | 45 | 60 | 75 |
|---|---|---|---|---|---|
| **Dendrite 1 (P13)** | -3.4(9.1) | -10.1(9.7) | -14.1(9.7) | -18.1(10.2) | -18.6(11.7) |
| **Dendrite 2 (P13)** | 3.2(9.3) | 3.4(9.5) | 1.5(10.3) | -1.0(11.2) | -4.0(11.0) |
| **Dendrite 3 (P13)** | -3.2(7.2) | -4.8(8.1) | -6.6(8.9) | -10.1(8.4) | -13.3(8.2) |
| **Dendrite 4 (P13)** | -0.6(8.3) | -2.7(8.6) | -4.5(10.0) | -7.0(9.3) | -11.6(9.0) |
| **Dendrite 5 (P13)** | -2.6(9.2) | -2.4(7.9) | -2.4(8.7) | -5.6(8.5) | -7.8(9.4) |
| **Dendrite 1 (P9)** | 3.2(7.7) | 1.1(10.7) | -1.1(12.3) | -2.3(14.4) | -4.4(14.7) |
| **Dendrite 2 (P9)** | -5.6(11.2) | -10.8(11.0) | -11.8(11.6) | -16.9(11.0) | -16.6(11.1) |
| **Dendrite 1 (P14)** | -7.0(11.4) | -12.1(11.8) | -15.7(12.3) | -19.2(10.9) | -20.3(12.2) |
| **Dendrite 2 (P14)** | -0.2(10.7) | -1.3(11.0) | -6.5(10.2) | -7.7(11.0) | -9.1(11.5) |
| **Dendrite 3 (P14)** | -0.8(11.6) | -1.5(10.4) | -4.6(10.6) | -5.7(10.0) | -7.8(11.1) |

| Number of removed spines | 90 | 105 | 120 | 135 | 150 |
|---|---|---|---|---|---|
| **Dendrite 1 (P13)** | -23.8(10.2) | -26.8(11.0) | -32.9(11.1) | -36.1(10.7) | -42.6(11.7) |
| **Dendrite 2 (P13)** | -8.7(10.1) | -8.5(10.2) | -12.6(10.5) | -15.4(10.8) | -19.8(9.4) |
| **Dendrite 3 (P13)** | -15.6(8.0) | -16.3(8.6) | -17.1(9.3) | -20.4(8.4) | -22.0(8.4) |
| **Dendrite 4 (P13)** | -10.9(9.1) | -15.0(10.2) | -16.4(8.7) | -16.6(9.3) | -18.7(9.3) |
| **Dendrite 5 (P13)** | -9.9(9.6) | -11.7(10.2) | -13.7(9.6) | -15.0(9.4) | -16.1(9.6) |
| **Dendrite 1 (P9)** | -3.6(14.4) | -9.4(14.7) | -11.3(14.6) | -18.5(12.5) | -20.0(16.0) |
| **Dendrite 2 (P9)** | -20.9(10.3) | -20.7(9.7) | -22.1(10.0) | -24.3(9.5) | -26.7(9.9) |
| **Dendrite 1 (P14)** | -23.0(11.5) | -27.1(10.5) | -30.8(9.6) | -32.4(9.7) | -35.3(9.9) |
| **Dendrite 2 (P14)** | -11.0(10.1) | -12.3(11.0) | -12.8(9.1) | -13.3(11.7) | -15.7(10.8) |
| **Dendrite 3 (P14)** | -9.8(11.5) | -12.5(10.0) | -12.2(9.9) | -15.5(9.8) | -16.0(10.1) |

**Supplementary Table S5: Percentage change of the grouping coefficient (GC) for all SomaTau- dendrites as a function of the number of removed spines.** The changes are shown in the cases when multiple spines are removed from the dendrites individually, in blocks of 3 and in blocks of 5 spines. The means are followed by standard deviations calculated over 100 trials.

| **Attack by removal of individual spines** | | | | | |
|---|---|---|---|---|---|
| **Number of removed spines** | **15** | **30** | **45** | **60** | **75** |
| **Dendrite 1 (P13)** | 0.1(0.9) | -0.2(1.4) | -0.1(1.8) | 0.0(2.0) | -0.4(2.0) |
| **Dendrite 2 (P13)** | 0.0(0.7) | 0.0(1.0) | 0.0(1.2) | 0.3(1.5) | -0.1(1.4) |
| **Dendrite 3 (P13)** | -0.1(0.3) | 0.0(0.4) | 0.0(0.5) | 0.0(0.6) | 0.1(0.8) |
| **Dendrite 4 (P13)** | 0.0(0.3) | 0.0(0.5) | 0.0(0.6) | -0.1(0.6) | 0.0(0.7) |
| **Dendrite 5 (P13)** | 0.0(0.3) | 0.0(0.4) | -0.1(0.5) | -0.1(0.6) | 0.0(0.6) |
| **Dendrite 1 (P9)** | -0.1(1.1) | -0.1(1.6) | 0.4(2.0) | 0.1(2.6) | 0.1(2.6) |
| **Dendrite 2 (P9)** | 0.0(0.2) | 0.0(0.3) | -0.1(0.4) | 0.0(0.5) | 0.0(0.6) |
| **Dendrite 1 (P14)** | 0.1(0.5) | 0.1(0.8) | 0.0(1.0) | -0.1(1.1) | -0.1(1.3) |
| **Dendrite 2 (P14)** | 0.0(0.3) | 0.0(0.4) | 0.1(0.5) | 0.0(0.6) | -0.1(0.6) |
| **Dendrite 3 (P14)** | 0.0(0.2) | 0.0(0.3) | 0.0(0.4) | 0.0(0.4) | 0.1(0.5) |
| **Number of removed spines** | **90** | **105** | **120** | **135** | **150** |
| **Dendrite 1 (P13)** | 0.2(2.5) | -0.1(3.0) | -0.2(3.0) | -0.1(4.1) | -0.1(4.4) |
| **Dendrite 2 (P13)** | 0.2(1.7) | 0.1(1.9) | 0.1(2.3) | 0.2(2.4) | 0.1(2.4) |
| **Dendrite 3 (P13)** | -0.1(0.7) | 0.0(1.0) | 0.0(1.0) | 0.0(1.2) | -0.1(1.1) |
| **Dendrite 4 (P13)** | 0.0(0.9) | -0.1(0.9) | -0.1(1.1) | 0.1(0.9) | 0.0(1.2) |
| **Dendrite 5 (P13)** | 0.0(0.7) | 0.0(0.7) | 0.0(0.8) | -0.2(0.9) | -0.1(1.0) |
| **Dendrite 1 (P9)** | -0.8(3.1) | 0.1(3.7) | -0.6(4.0) | -0.2(5.1) | -0.2(5.0) |
| **Dendrite 2 (P9)** | 0.0(0.7) | 0.0(0.7) | 0.0(0.8) | 0.0(0.9) | 0.0(0.9) |
| **Dendrite 1 (P14)** | 0.1(1.3) | 0.2(1.7) | 0.0(1.7) | 0.4(2.0) | -0.1(2.2) |
| **Dendrite 2 (P14)** | 0.1(0.7) | 0.0(0.9) | 0.0(0.8) | 0.0(1.0) | -0.1(1.0) |
| **Dendrite 3 (P14)** | 0.0(0.5) | 0.0(0.6) | 0.1(0.6) | 0.0(0.7) | 0.0(0.7) |
| **Attack by removal of spines in blocks of 3** | | | | | |
| **Number of removed spines** | **15** | **30** | **45** | **60** | **75** |
| **Dendrite 1 (P13)** | 0.4(1.4) | 1.5(2.3) | 3.7(3.3) | 5.8(4.7) | 8.9(5.5) |
| **Dendrite 2 (P13)** | 0.5(1.1) | 0.8(1.5) | 1.7(2.4) | 2.3(2.4) | 3.9(3.2) |
| **Dendrite 3 (P13)** | 0.2(0.5) | 0.3(0.7) | 0.9(1.1) | 1.2(1.3) | 2.2(1.5) |
| **Dendrite 4 (P13)** | 0.2(0.6) | 0.4(0.8) | 0.9(1.0) | 1.4(1.2) | 2.1(1.5) |
| **Dendrite 5 (P13)** | 0.2(0.5) | 0.4(0.7) | 0.6(0.9) | 1.0(1.1) | 1.3(1.1) |
| **Dendrite 1 (P9)** | 0.8(1.7) | 2.1(2.6) | 3.4(3.5) | 5.8(4.9) | 9.7(5.2) |
| **Dendrite 2 (P9)** | 0.2(0.4) | 0.4(0.7) | 0.7(0.8) | 1.4(1.0) | 2.1(1.4) |
| **Dendrite 1 (P14)** | 0.4(1.1) | 0.8(1.3) | 1.5(1.7) | 2.6(2.5) | 4.1(3.1) |
| **Dendrite 2 (P14)** | 0.2(0.5) | 0.4(0.7) | 0.6(0.8) | 1.1(1.1) | 1.3(1.2) |
| **Dendrite 3 (P14)** | 0.2(0.4) | 0.3(0.5) | 0.4(0.6) | 0.7(0.7) | 1.1(0.9) |

| Number of removed spines | 90 | 105 | 120 | 135 | 150 |
|---|---|---|---|---|---|
| **Dendrite 1 (P13)** | 12.3(6.9) | 20.7(8.9) | 27.4(13.4) | 39.2(15.0) | 61.4(22.5) |
| **Dendrite 2 (P13)** | 5.4(3.1) | 6.6(4.6) | 8.9(4.9) | 10.8(5.8) | 13.7(6.4) |
| **Dendrite 3 (P13)** | 3.0(1.7) | 3.6(1.9) | 5.1(2.4) | 6.3(2.6) | 7.4(2.6) |
| **Dendrite 4 (P13)** | 2.7(1.7) | 3.5(1.9) | 4.5(2.2) | 5.9(2.4) | 7.5(3.1) |
| **Dendrite 5 (P13)** | 2.0(1.5) | 2.5(1.7) | 3.7(2.1) | 4.4(2.1) | 4.9(2.1) |
| **Dendrite 1 (P9)** | 14.1(6.6) | 18.3(9.5) | 25.9(10.5) | 36.6(12.3) | 42.1(9.2) |
| **Dendrite 2 (P9)** | 2.5(1.4) | 3.7(2.0) | 4.9(2.0) | 6.4(2.5) | 7.4(3.1) |
| **Dendrite 1 (P14)** | 5.3(3.2) | 6.8(4.3) | 10.8(4.1) | 12.2(5.0) | 16.5(6.0) |
| **Dendrite 2 (P14)** | 2.0(1.5) | 2.2(1.7) | 3.6(1.7) | 4.1(2.2) | 5.1(2.3) |
| **Dendrite 3 (P14)** | 1.3(1.0) | 1.6(1.2) | 2.2(1.4) | 2.9(1.3) | 3.4(1.5) |
| **Attack by removal of spines in blocks of 5** | | | | | |
| **Number of removed spines** | **15** | **30** | **45** | **60** | **75** |
| **Dendrite 1 (P13)** | 1.2(1.9) | 2.2(3.1) | 4.7(3.8) | 7.0(5.1) | 9.8(5.9) |
| **Dendrite 2 (P13)** | 0.6(1.4) | 0.8(2.1) | 1.8(2.7) | 2.7(3.3) | 4.0(3.3) |
| **Dendrite 3 (P13)** | 0.1(0.6) | 0.7(0.9) | 1.2(1.3) | 1.7(1.6) | 2.9(2.0) |
| **Dendrite 4 (P13)** | 0.3(0.7) | 1.0(1.1) | 1.1(1.6) | 2.1(1.8) | 2.7(2.2) |
| **Dendrite 5 (P13)** | 0.2(0.5) | 0.5(0.9) | 0.8(1.2) | 1.4(1.3) | 1.9(1.7) |
| **Dendrite 1 (P9)** | 1.3(2.2) | 3.5(3.3) | 5.5(4.7) | 6.9(6.5) | 12.4(6.7) |
| **Dendrite 2 (P9)** | 0.4(0.5) | 0.8(0.9) | 1.3(1.1) | 1.9(1.3) | 2.7(1.5) |
| **Dendrite 1 (P14)** | 0.7(1.4) | 1.0(1.7) | 1.9(2.2) | 3.1(3.0) | 4.9(3.3) |
| **Dendrite 2 (P14)** | 0.2(0.6) | 0.5(1.0) | 0.7(1.0) | 1.5(1.3) | 1.8(1.6) |
| **Dendrite 3 (P14)** | 0.2(0.4) | 0.4(0.6) | 0.7(0.8) | 1.0(1.0) | 1.2(1.1) |
| **Number of removed spines** | **90** | **105** | **120** | **135** | **150** |
| **Dendrite 1 (P13)** | 14.5(8.2) | 23.9(12.1) | 32.9(14.3) | 42.9(22.9) | 69.3(30.9) |
| **Dendrite 2 (P13)** | 5.8(4.2) | 7.4(5.4) | 9.6(5.5) | 12.2(6.4) | 16.0(8.2) |
| **Dendrite 3 (P13)** | 3.5(2.5) | 4.8(2.6) | 5.6(2.8) | 7.4(3.7) | 9.4(3.9) |
| **Dendrite 4 (P13)** | 3.3(2.2) | 4.3(2.7) | 5.8(2.6) | 7.1(3.6) | 8.3(3.3) |
| **Dendrite 5 (P13)** | 2.3(1.8) | 3.2(1.8) | 4.3(2.2) | 5.4(3.0) | 6.8(2.7) |
| **Dendrite 1 (P9)** | 16.0(7.8) | 21.0(10.5) | 28.9(12.0) | 39.7(12.9) | 49.6(14.3) |
| **Dendrite 2 (P9)** | 3.8(1.8) | 4.8(2.1) | 5.9(2.4) | 7.9(3.4) | 8.8(3.0) |
| **Dendrite 1 (P14)** | 6.3(3.9) | 8.9(4.5) | 10.4(5.4) | 14.2(5.6) | 18.6(7.1) |
| **Dendrite 2 (P14)** | 2.7(1.7) | 3.5(2.1) | 4.2(2.3) | 5.0(2.5) | 5.7(2.9) |
| **Dendrite 3 (P14)** | 1.8(1.5) | 2.1(1.4) | 2.7(1.6) | 3.7(1.7) | 4.6(2.2) |

### Replication in two alternative methods for construction of graphs

The results shown in the main text are consistent with those obtained using alternative methods to construct the dendrite-specific graphs. First, we build the graphs by estimating the strength of association between a pair of spines as a fraction of the total length of the corresponding dendrite. In particular, denoting the distance between spines $i$ and $k$ as $d_{ik}$, we calculate the strength of association between the two spines as

$$w_{ik}{}^{1} = \frac{1}{d_{ik}/L} = \frac{L}{d_{ik}},$$

where $L$ is the total length of the dendrite expressed in μm.

In the second alternative approach, we calculate the association between the two spines as

$$w_{ik}{}^{2} = \ln(d_{ik}).$$

Since taking the logarithm of the distances results in negative numbers, and the graph measures do not have clear extensions for negative weights, we add a constant to all weights and analyze the resulting graphs. Specifically,

$$w_{ik}{}^{2} = \ln(\mathrm{d_{min}}) + \ln(d_{ik}),$$

where $\mathrm{d_{min}}$ denotes the minimum distance of any two spines across all dendrites.

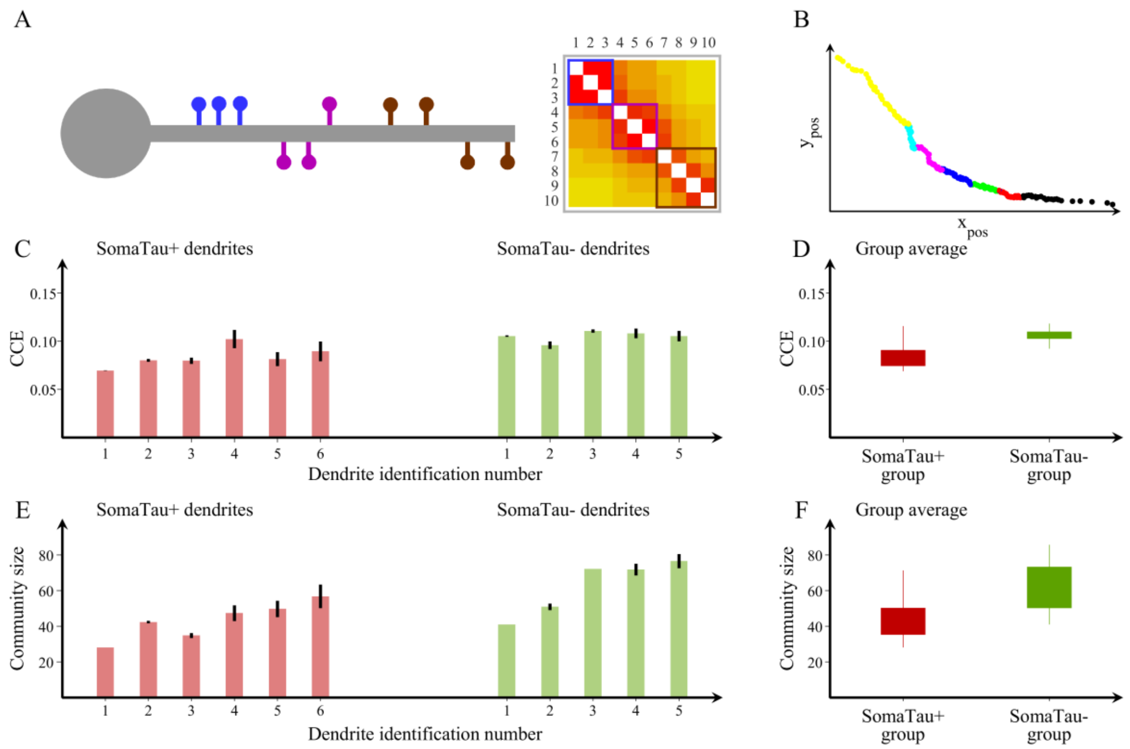

**Supplementary Figure S5. Spine organization into communities for individual P13 – graphs calculated by alternative method 1.**

(**A**) Schematic representation of a dendrite with spines that can be clearly separated into three distinct communities and their corresponding adjacency matrix showing the blocks of connections that correspond to each community. (**B**) We also show an example of a community structure for one of the dendrites assessed in the current study. The community structure is assessed using the average distance between spines that belong to each community (characteristic community extension, CCE) and the number of spines in each community. $Y_{pos}$ and $X_{pos}$, (μm). (**C, E**) We show the values obtained in these two measures for each SomaTau+ (red, n= 6) and SomaTau- (green, n = 5) dendrite, which are computed by calculating the community structure over 100 trials. In addition, we include (**D**) boxplots with the group averages for the CCE and (**F**) the average community size or number of spines for each community, which are both smaller in SomaTau+ compared to SomaTau- dendrites. The bottom and the top edges of the boxplots denote the $25^{th}$ and $75^{th}$ percentiles of the data, respectively. The whiskers extend to the largest and smallest data points. The results are similar after excluding the outlier.

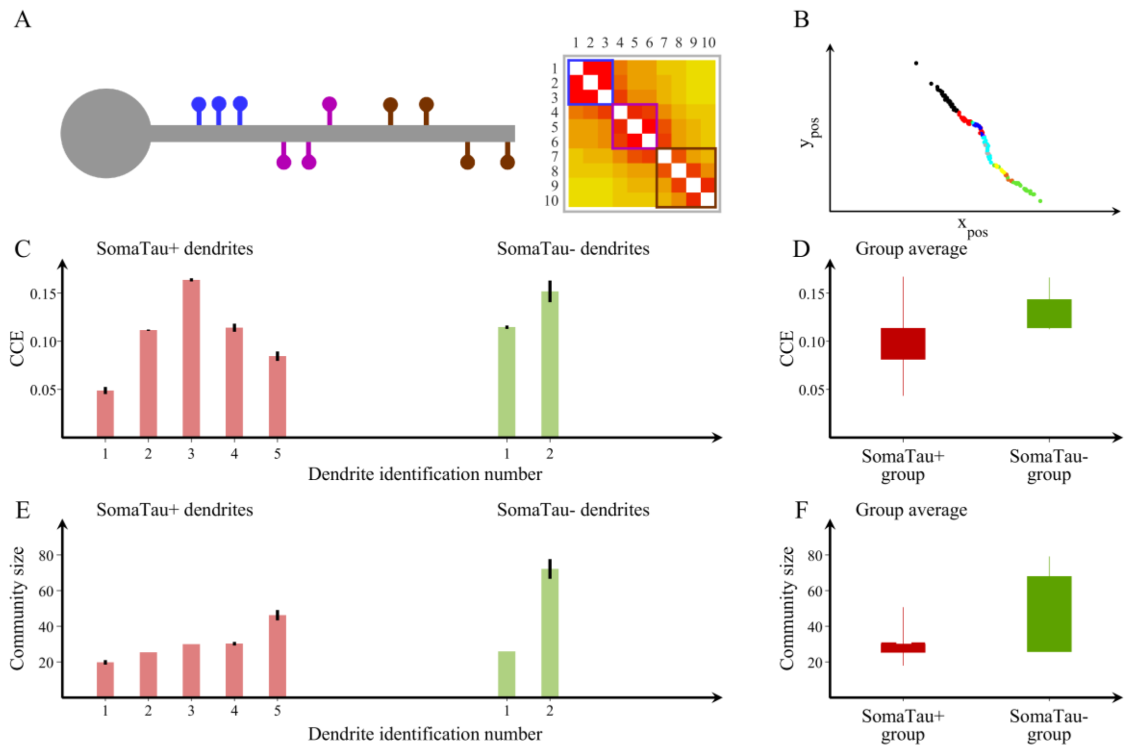

**Supplementary Figure S6. Spine organization into communities for individual P9 – graphs calculated by alternative method 1.**

(**A**) Schematic representation of a dendrite with spines that can be clearly separated into three distinct communities and their corresponding adjacency matrix showing the blocks of connections that correspond to each community. (**B**) We also show an example of a community structure for one of the dendrites assessed in the current study. The community structure is assessed using the average distance between spines that belong to each community (characteristic community extension, CCE) and the number of spines in each community. $Y_{pos}$ and $X_{pos}$, (μm). (**C, E**) We show the values obtained in these two measures for each SomaTau+ (red, n= 5) and SomaTau- (green, n = 2) dendrite, which are computed by calculating the community structure over 100 trials. In addition, we include (**D**) boxplots with the group averages for the CCE and (**F**) the average community size or number of spines for each community, which are both smaller in SomaTau+ compared to SomaTau- dendrites. The bottom and the top edges of the boxplots denote the 25th and 75th percentiles of the data, respectively. The whiskers extend to the largest and smallest data points. The results are similar after excluding the outlier.

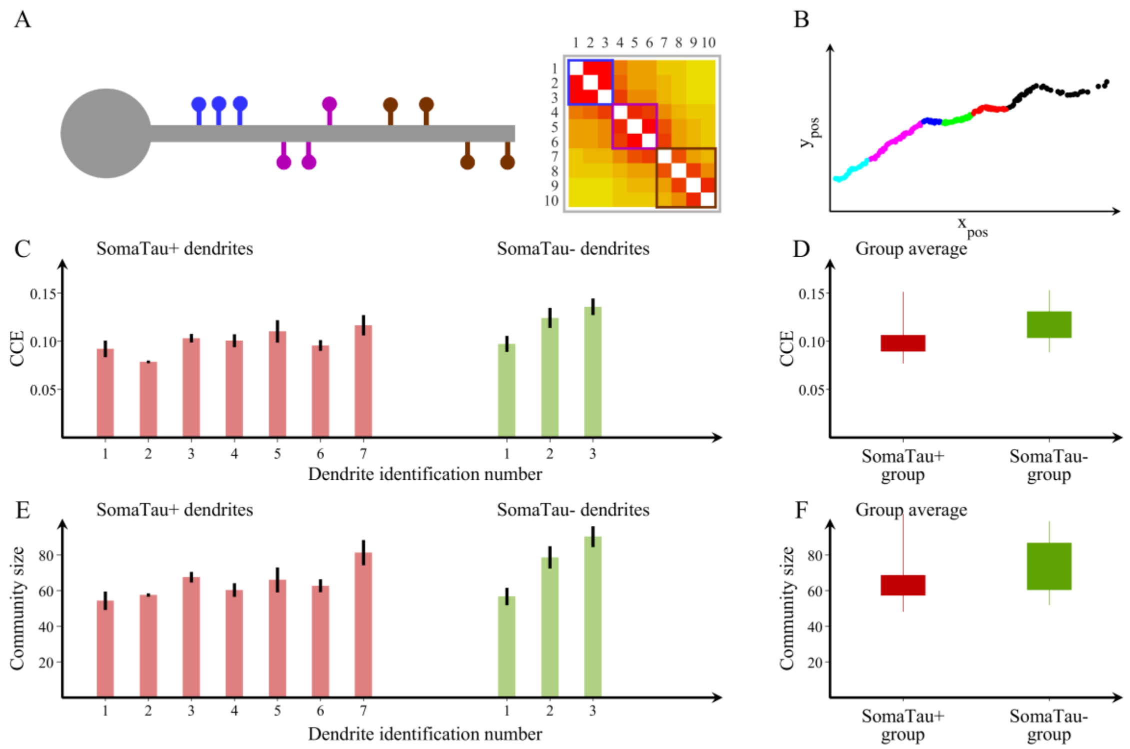

**Supplementary Figure S7. Spine organization into communities for individual P14 – graphs calculated by alternative method 1.**

(**A**) Schematic representation of a dendrite with spines that can be clearly separated into three distinct communities and their corresponding adjacency matrix showing the blocks of connections that correspond to each community. (**B**) We also show an example of a community structure for one of the dendrites assessed in the current study. The community structure is assessed using the average distance between spines that belong to each community (characteristic community extension, CCE) and the number of spines in each community. $Y_{pos}$ and $X_{pos}$, (μm). (**C, E**) We show the values obtained in these two measures for each SomaTau+ (red, n= 7) and SomaTau- (green, n = 3) dendrite, which are computed by calculating the community structure over 100 trials. In addition, we include (**D**) boxplots with the group averages for the CCE and (**F**) the average community size or number of spines for each community, which are both smaller in SomaTau+ compared to SomaTau- dendrites. The bottom and the top edges of the boxplots denote the 25$^{th}$ and 75$^{th}$ percentiles of the data, respectively. The whiskers extend to the largest and smallest data points. The results are similar after excluding the outlier.

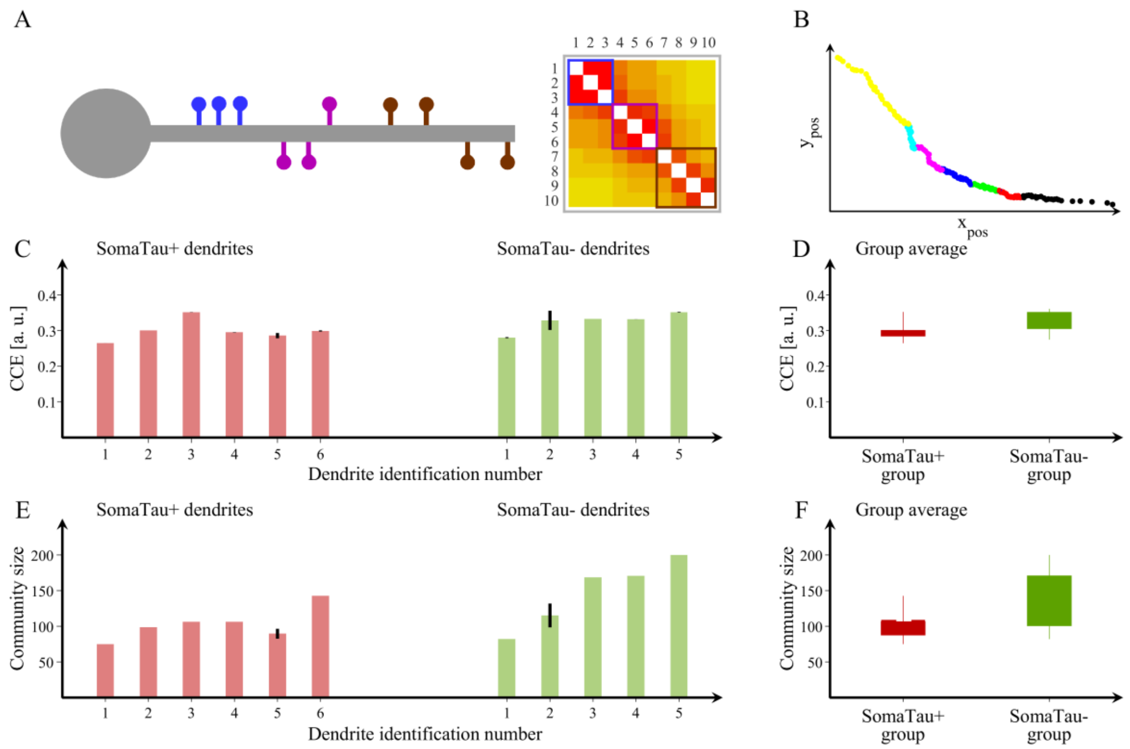

**Supplementary Figure S8. Spine organization into communities for individual P13 – graphs calculated by alternative method 2.**

(**A**) Schematic representation of a dendrite with spines that can be clearly separated into three distinct communities and their corresponding adjacency matrix showing the blocks of connections that correspond to each community. (**B**) We also show an example of a community structure for one of the dendrites assessed in the current study. The community structure is assessed using the average distance between spines that belong to each community (characteristic community extension, CCE) and the number of spines in each community. $Y_{pos}$ and $X_{pos}$, (μm). (**C, E**) We show the values obtained in these two measures for each SomaTau+ (red, n= 6) and SomaTau- (green, n = 5) dendrite, which are computed by calculating the community structure over 100 trials. In addition, we include (**D**) boxplots with the group averages for the CCE and (**F**) the average community size or number of spines for each community, which are both smaller in SomaTau+ compared to SomaTau- dendrites. The bottom and the top edges of the boxplots denote the 25$^{th}$ and 75$^{th}$ percentiles of the data, respectively. The whiskers extend to the largest and smallest data points. The results are similar after excluding the outlier.

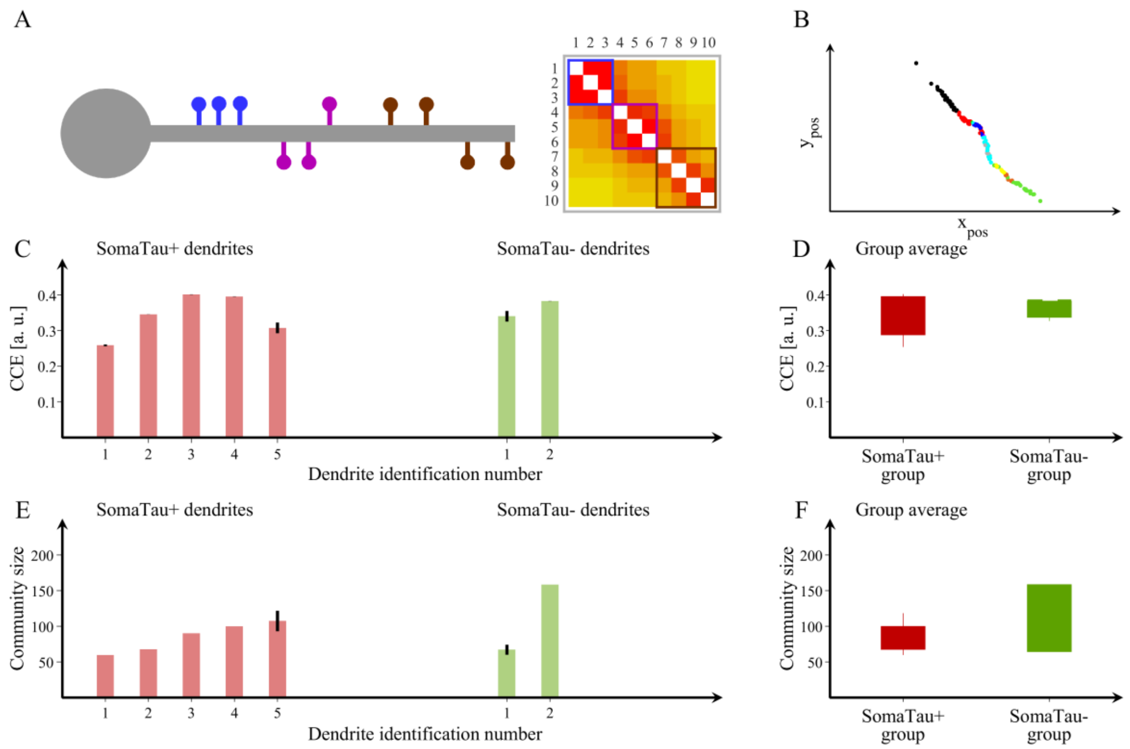

**Supplementary Figure S9. Spine organization into communities for individual P9 – graphs calculated by alternative method 2.**

(**A**) Schematic representation of a dendrite with spines that can be clearly separated into three distinct communities and their corresponding adjacency matrix showing the blocks of connections that correspond to each community. (**B**) We also show an example of a community structure for one of the dendrites assessed in the current study. The community structure is assessed using the average distance between spines that belong to each community (characteristic community extension, CCE) and the number of spines in each community. $Y_{pos}$ and $X_{pos}$, (μm). (**C, E**) We show the values obtained in these two measures for each SomaTau+ (red, n= 5) and SomaTau- (green, n = 2) dendrite, which are computed by calculating the community structure over 100 trials. In addition, we include (**D**) boxplots with the group averages for the CCE and (**F**) the average community size or number of spines for each community, which are both smaller in SomaTau+ compared to SomaTau- dendrites. The bottom and the top edges of the boxplots denote the 25$^{th}$ and 75$^{th}$ percentiles of the data, respectively. The whiskers extend to the largest and smallest data points. The results are similar after excluding the outlier.

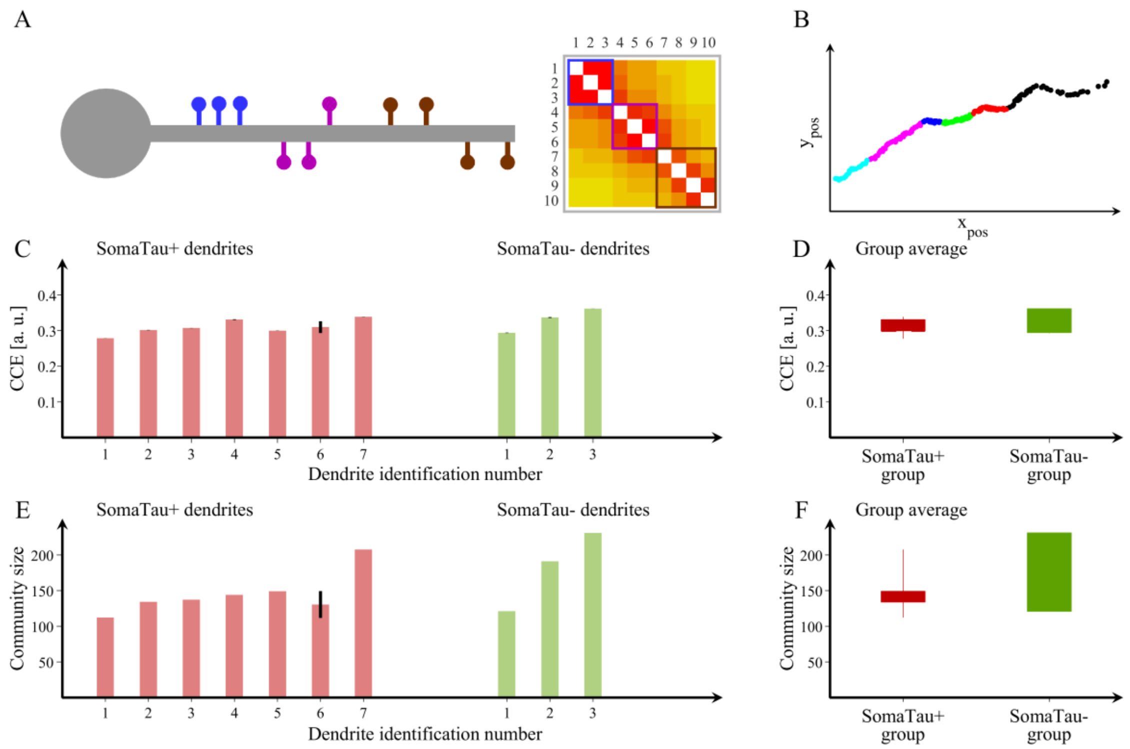

**Supplementary Figure S10. Spine organization into communities for individual P14 – graphs calculated by alternative method 2.**

(**A**) Schematic representation of a dendrite with spines that can be clearly separated into three distinct communities and their corresponding adjacency matrix showing the blocks of connections that correspond to each community. (**B**) We also show an example of a community structure for one of the dendrites assessed in the current study. The community structure is assessed using the average distance between spines that belong to each community (characteristic community extension, CCE) and the number of spines in each community. $Y_{pos}$ and $X_{pos}$, (μm). (**C, E**) We show the values obtained in these two measures for each SomaTau+ (red, n= 7) and SomaTau- (green, n = 3) dendrite, which are computed by calculating the community structure over 100 trials. In addition, we include (**D**) boxplots with the group averages for the CCE and (**F**) the average community size or number of spines for each community, which are both smaller in SomaTau+ compared to SomaTau- dendrites. The bottom and the top edges of the boxplots denote the 25$^{th}$ and 75$^{th}$ percentiles of the data, respectively. The whiskers extend to the largest and smallest data points. The results are similar after excluding the outlier.

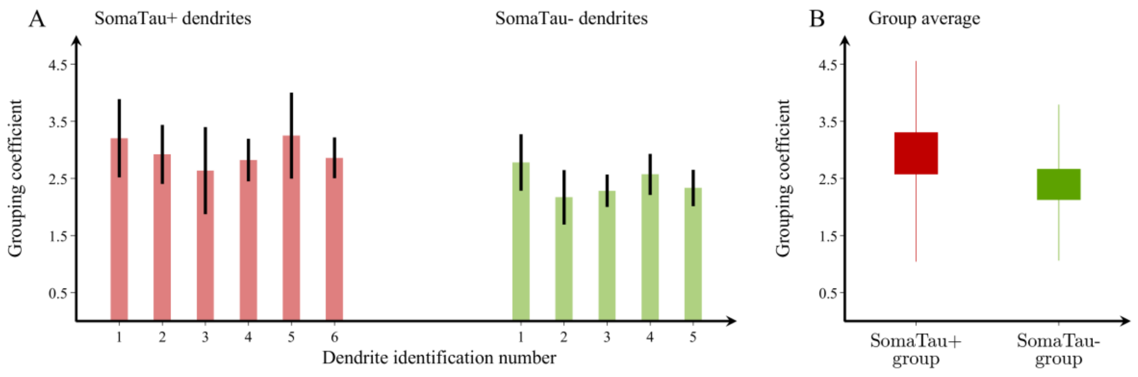

**Supplementary Figure S11. Mean grouping coefficient in dendrites with and without tau pathology for individual P13 – graphs calculated by alternative method 1.**

(**A**) Mean grouping coefficient in each single SomaTau+ (n = 6; red) and SomaTau- (n = 5; green) dendrite. Boxplots with the mean grouping coefficients in the SomaTau- and SomaTau+ dendrites. (**B**) The permutation analyses show a higher mean grouping coefficient in the Tau+ compared to the Tau- group ($p < 0.001$). In all boxplots, their bottom and the top edges denote the 25$^{th}$ and 75$^{th}$ percentiles of the data, respectively. The whiskers extend to the largest and smallest data points. The results are similar after excluding the outliers.

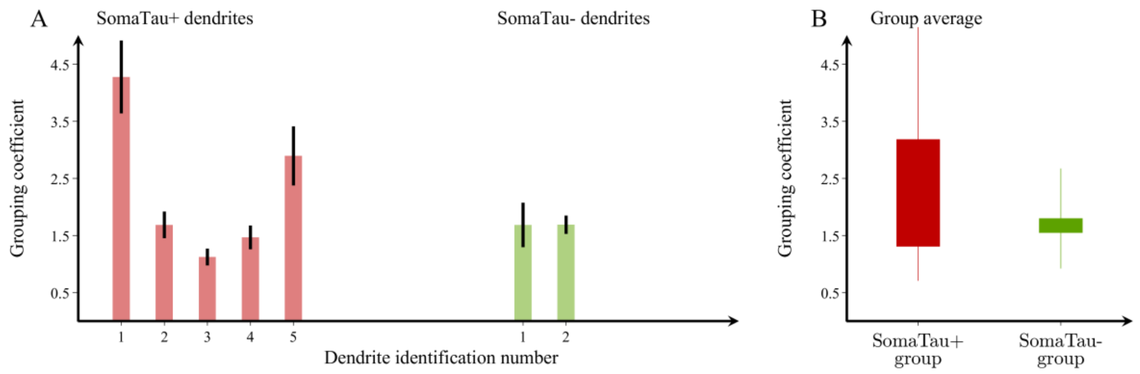

**Supplementary Figure S12. Mean grouping coefficient in dendrites with and without tau pathology for individual P9 – graphs calculated by alternative method 1.**

(**A**) Mean grouping coefficient in each single SomaTau+ (n = 5; red) and SomaTau- (n = 2; green) dendrite. Boxplots with the mean grouping coefficients in the SomaTau- and SomaTau+ dendrites. (**B**) The permutation analyses show a higher mean grouping coefficient in the Tau+ compared to the Tau- group ($p < 0.001$). In all boxplots, their bottom and the top edges denote the 25$^{th}$ and 75$^{th}$ percentiles of the data, respectively. The whiskers extend to the largest and smallest data points. The results are similar after excluding the outliers.

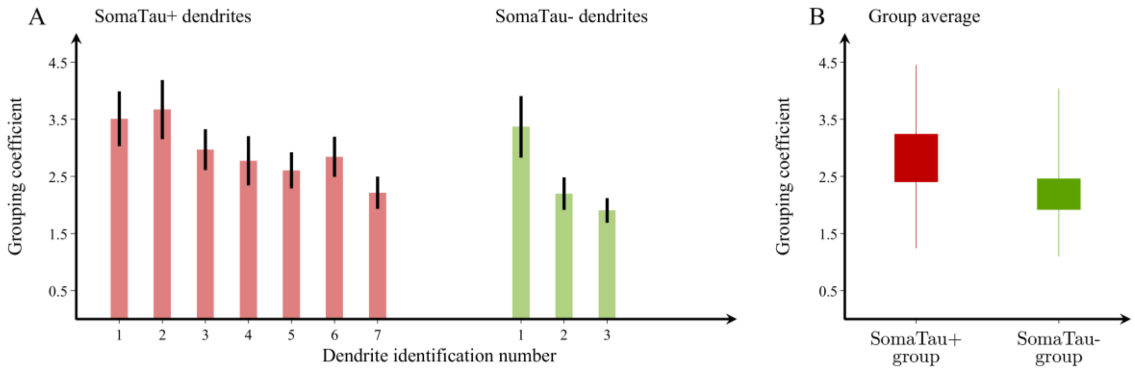

**Supplementary Figure S13. Mean grouping coefficient in dendrites with and without tau pathology for individual P14 – graphs calculated by alternative method 1.**

(**A**) Mean grouping coefficient in each single SomaTau+ (n = 7; red) and SomaTau- (n = 3; green) dendrite. Boxplots with the mean grouping coefficients in the SomaTau- and SomaTau+ dendrites. (**B**) The permutation analyses show a higher mean grouping coefficient in the Tau+ compared to the Tau- group ($p < 0.001$). In all boxplots, their bottom and the top edges denote the 25$^{th}$ and 75$^{th}$ percentiles of the data, respectively. The whiskers extend to the largest and smallest data points. The results are similar after excluding the outliers.

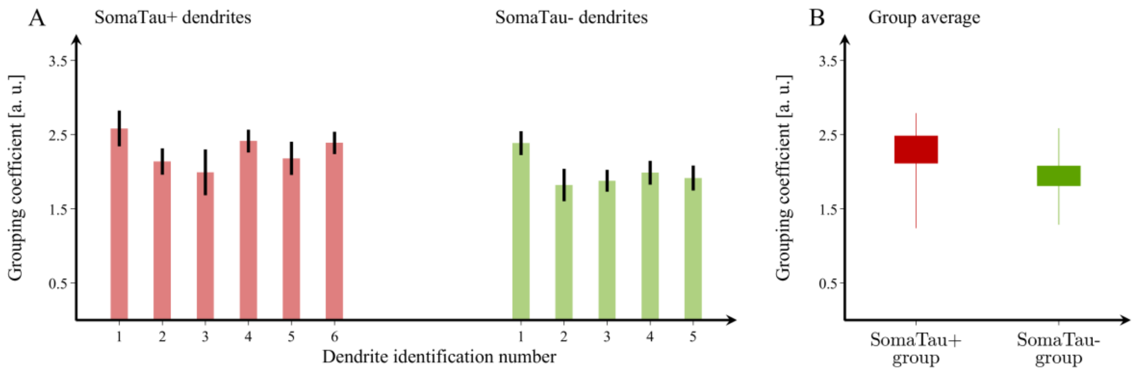

**Supplementary Figure S14. Mean grouping coefficient in dendrites with and without tau pathology for individual P13 – graphs calculated by alternative method 2.**

(**A**) Mean grouping coefficient in each single SomaTau+ (n = 6; red) and SomaTau- (n = 5; green) dendrite. Boxplots with the mean grouping coefficients in the SomaTau- and SomaTau+ dendrites. (**B**) The permutation analyses show a higher mean grouping coefficient in the Tau+ compared to the Tau- group ($p < 0.001$). In all boxplots, their bottom and the top edges denote the $25^{th}$ and $75^{th}$ percentiles of the data, respectively. The whiskers extend to the largest and smallest data points. The results are similar after excluding the outliers.

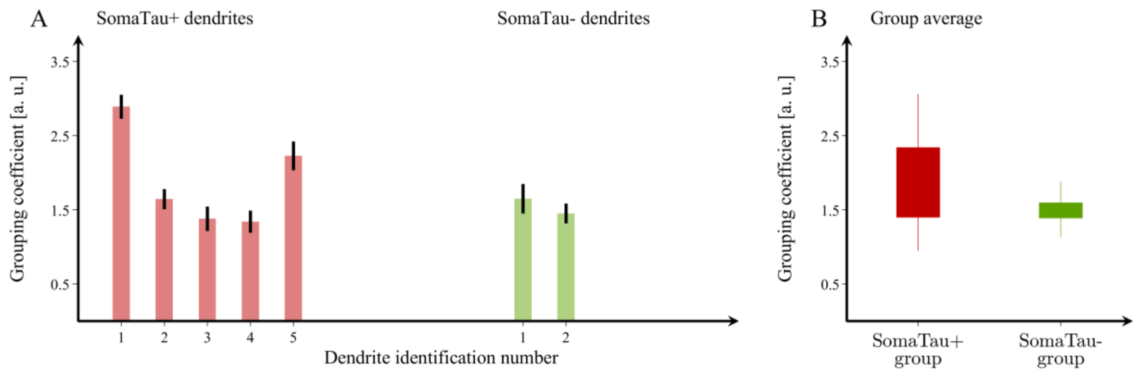

**Supplementary Figure S15. Mean grouping coefficient in dendrites with and without tau pathology for individual P9 – graphs calculated by alternative method 2.**

(**A**) Mean grouping coefficient in each single SomaTau+ (n = 5; red) and SomaTau- (n = 2; green) dendrite. Boxplots with the mean grouping coefficients in the SomaTau- and SomaTau+ dendrites. (**B**) The permutation analyses show a higher mean grouping coefficient in the Tau+ compared to the Tau- group ($p < 0.001$). In all boxplots, their bottom and the top edges denote the 25$^{th}$ and 75$^{th}$ percentiles of the data, respectively. The whiskers extend to the largest and smallest data points. The results are similar after excluding the outliers.

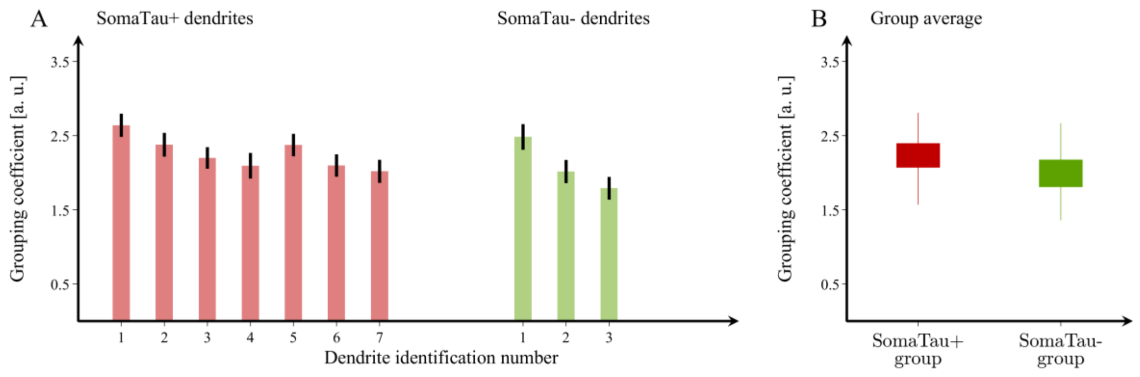

**Supplementary Figure S16. Mean grouping coefficient in dendrites with and without tau pathology for individual P14 – graphs calculated by alternative method 2.**

(**A**) Mean grouping coefficient in each single SomaTau+ (n = 7; red) and SomaTau- (n = 3; green) dendrite. Boxplots with the mean grouping coefficients in the SomaTau- and SomaTau+ dendrites. (**B**) The permutation analyses show a higher mean grouping coefficient in the Tau+ compared to the Tau- group ($p < 0.001$). In all boxplots, their bottom and the top edges denote the $25^{th}$ and $75^{th}$ percentiles of the data, respectively. The whiskers extend to the largest and smallest data points. The results are similar after excluding the outliers.

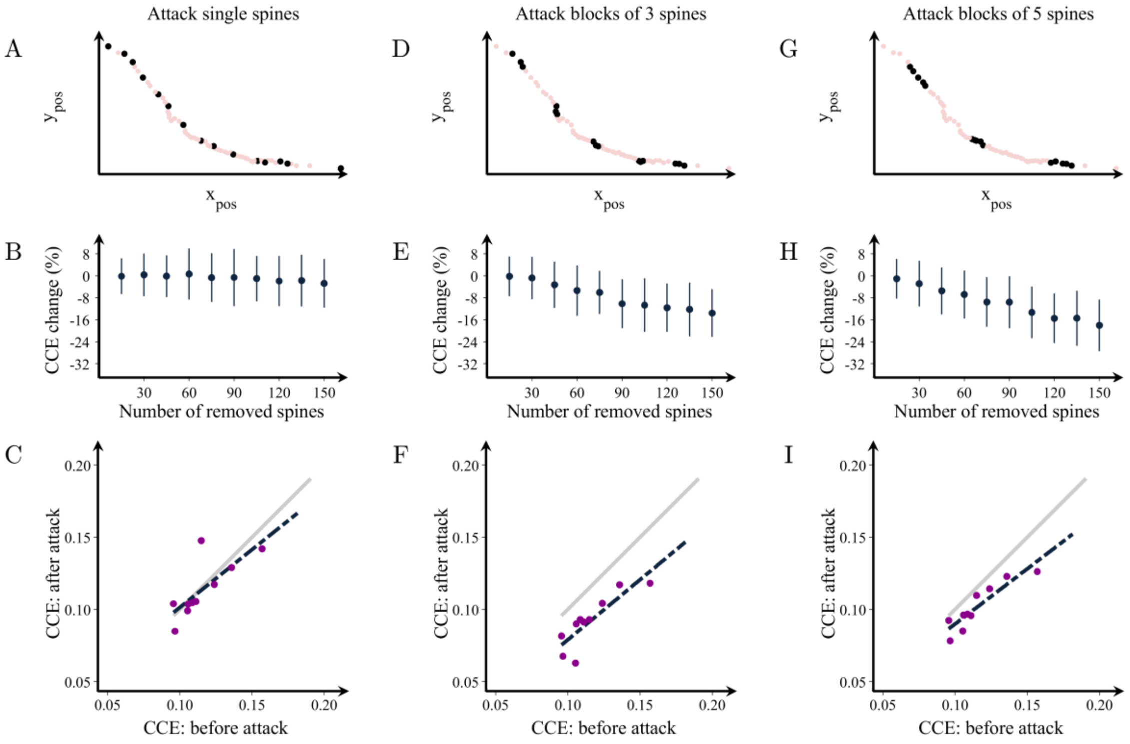

**Supplementary Figure S17. Changes in the organization of dendritic spines after attacks - graphs calculated by alternative method 1.** Examples of attacks on spines in a representative dendrite at (**A**) random locations or in clusters of (**D**) 3 and (**G**) 5 spines. (**B, E, H**) Percentage change in the characteristic community extension (CCE) as a function of the number of removed spines in the three cases, respectively. CCE in the attacked vs. the non-attacked dendrites after the removal of 150 spines in groups of (**C**) 1, (**F**) 3 and (**I**) 5. The grey line shows the theoretical line of no change in the CCE, while the black line is the line that best fits the observed attack data. Means and standard deviations are computed over 100 random attacks.

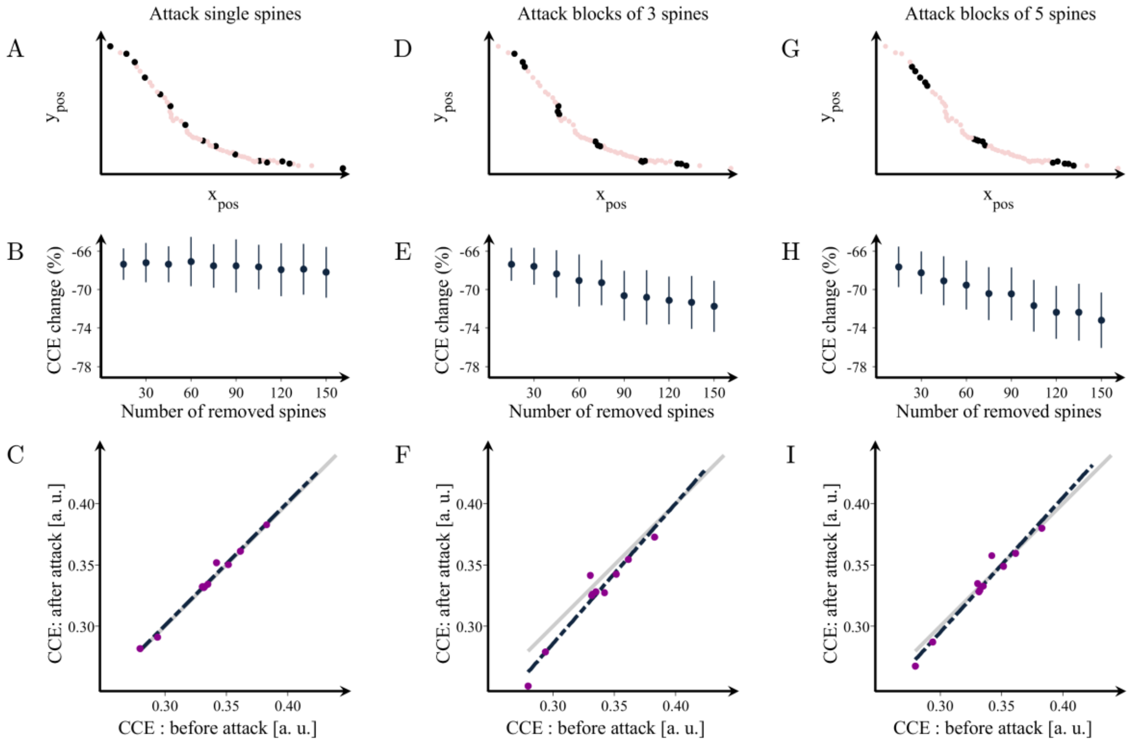

**Supplementary Figure S18. Changes in the organization of dendritic spines after attacks - graphs calculated by alternative method 2.** Examples of attacks on spines in a representative dendrite at (**A**) random locations or in clusters of (**D**) 3 and (**G**) 5 spines. (**B, E, H**) Percentage change in the characteristic community extension (CCE) as a function of the number of removed spines in the three cases, respectively. CCE in the attacked vs. the non-attacked dendrites after the removal of 150 spines in groups of (**C**) 1, (**F**) 3 and (**I**) 5. The grey line shows the theoretical line of no change in the CCE, while the black line is the line that best fits the observed attack data. Means and standard deviations are computed over 100 random attacks.

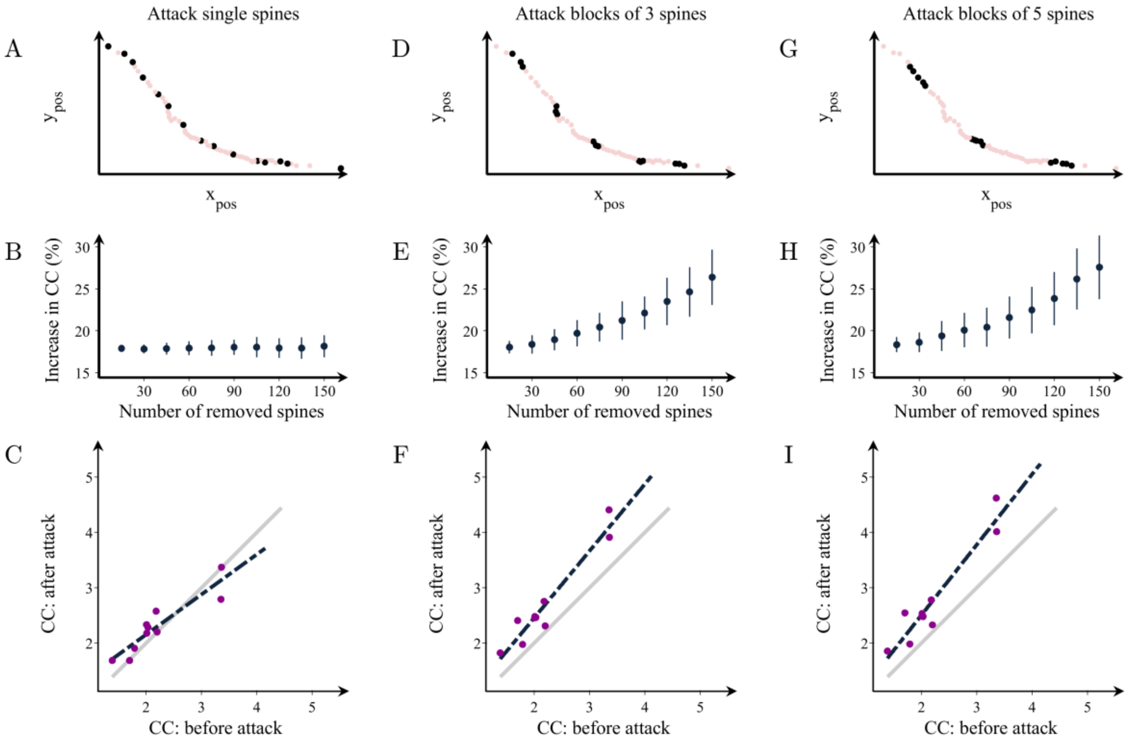

**Supplementary Figure S19. Changes in the mean grouping coefficient after random attacks of spines in healthy dendrites - graphs calculated by alternative method 1.** Illustration of attacks on random spines in a representative dendrite in groups of (**A**) 1, (**D**) 3 and (**G**) 5. (**B, E, H**) Percentage increases in the mean grouping coefficient (GC) in that dendrite as a function of the number of removed spines in the three cases respectively. Mean grouping coefficient of the attacked vs. healthy dendrites after the removal of 150 spines in groups of (**C**) 1, (**F**) 3 and (**I**) 5. The grey line shows the theoretical line of no change in the grouping coefficient, while the black line is the line that best fits the observed data after the attacks. Means and standard deviations are computed over 100 random attacks.

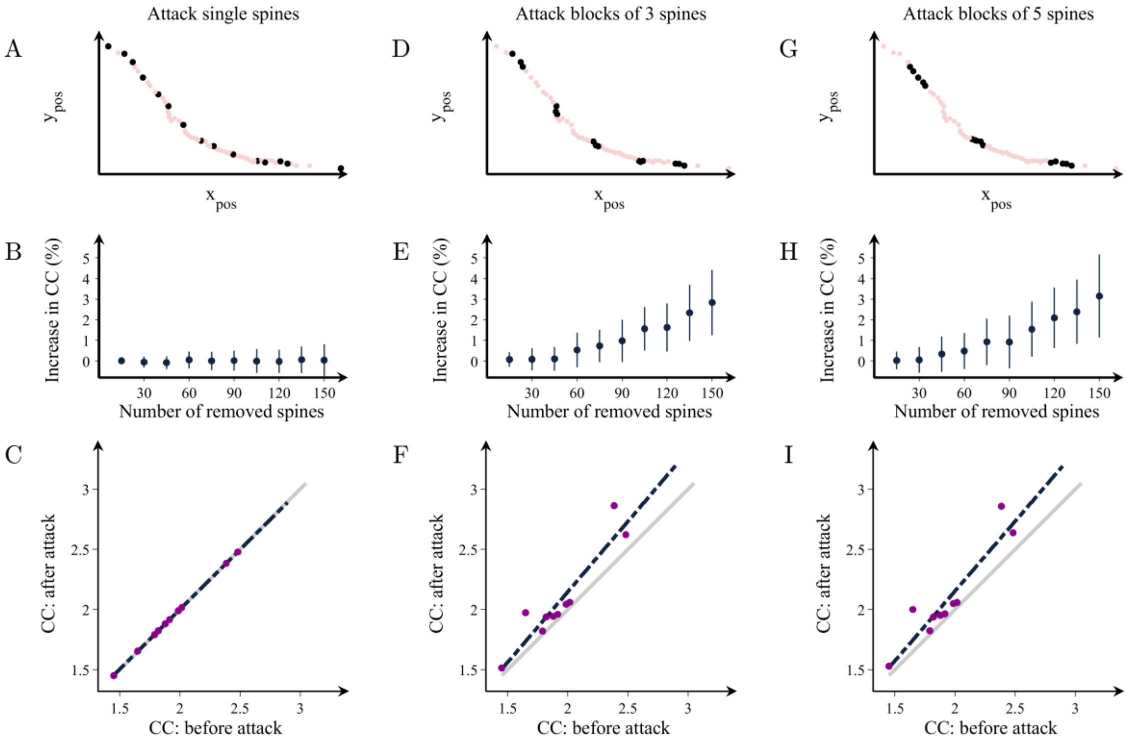

**Supplementary Figure S20. Changes in the mean grouping coefficient after random attacks of spines in healthy dendrites - graphs calculated by alternative method 2.** Illustration of attacks on random spines in a representative dendrite in groups of (**A**) 1, (**D**) 3 and (**G**) 5. (**B, E, H**) Percentage increases in the mean grouping coefficient (GC) in that dendrite as a function of the number of removed spines in the three cases respectively. Mean grouping coefficient of the attacked vs. healthy dendrites after the removal of 150 spines in groups of (**C**) 1, (**F**) 3 and (**I**) 5. The grey line shows the theoretical line of no change in the grouping coefficient, while the black line is the line that best fits the observed data after the attacks. Means and standard deviations are computed over 100 random attacks.

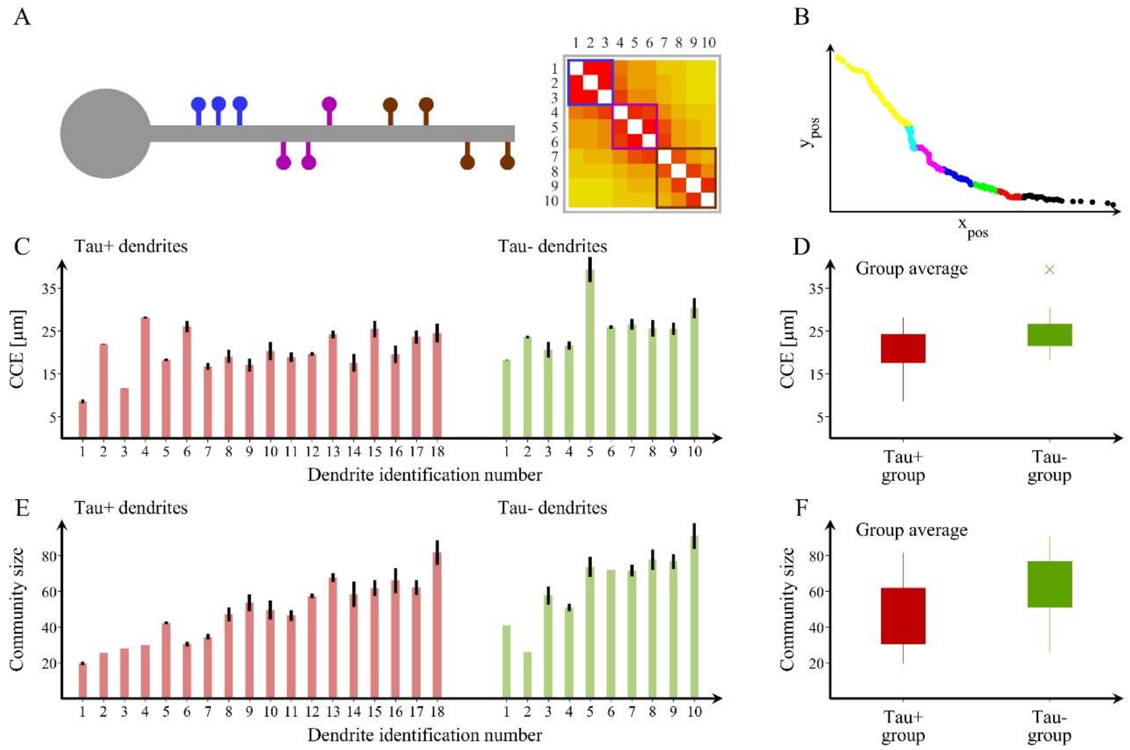

**Supplementary Figure 21. Spine organization into communities.**

(**A**) Schematic representation of a dendrite with spines that can be clearly separated into three distinct communities and their corresponding adjacency matrix showing the blocks of connections that correspond to each community. (**B**) We also show an example of a community structure for one of the dendrites assessed in the current study with seven communities. The community structure is assessed using the average distance between spines that belong to each community (characteristic community extension, CCE) and the number of spines in each community. $Y_{pos}$ and $X_{pos}$, (µm). (**C, E**) We show the values obtained in these two measures for each Tau+ (red, n= 18) and Tau- (green, n = 10) dendrite, which are computed by calculating the community structure over 100 trials. In addition, we include (**D**) boxplots with the group averages for the CCE and (**F**) the average community size or number of spines for each community, which are both smaller in Tau+ compared to Tau- dendrites. The bottom and the top edges of the boxplots denote the 25$^{th}$ and 75$^{th}$ percentiles of the data, respectively. The whiskers extend to the largest and smallest data points that are not considered outliers. The outliers are plotted with the “x” symbol. The results are similar after excluding the outlier.

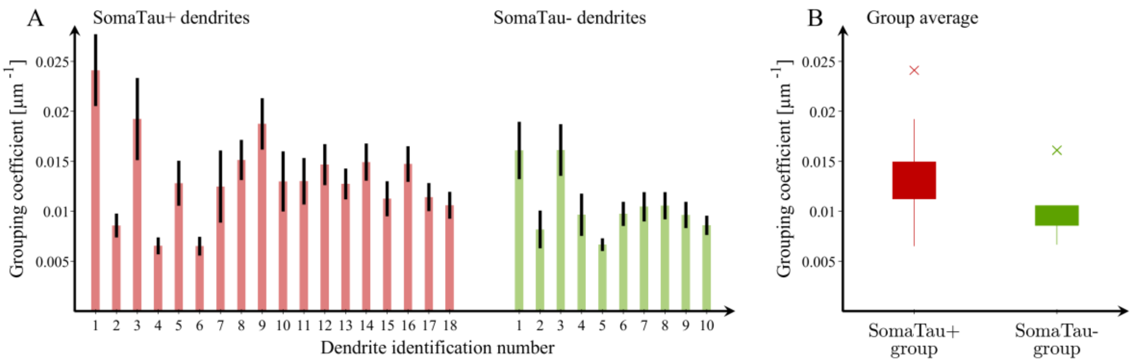

**Supplementary Figure 22. Mean grouping coefficient in dendrites with and without tau pathology.**

(**A**) Mean grouping coefficient in each individual Tau+ (n = 18; red) and Tau- (n = 10; green) dendrite. Boxplots with the mean grouping coefficients in the Tau- and Tau+ dendrites. (**B**) The permutation analyses show a higher mean grouping coefficient in the Tau+ compared to the Tau- group ($p < 0.001$). In all boxplots, their bottom and the top edges denote the 25$^{th}$ and 75$^{th}$ percentiles of the data, respectively. The whiskers extend to the largest and smallest data points that are not considered outliers. The outliers are plotted with the “x” symbol. The results are similar after excluding the outliers.
